# NQO1 phase condensation promotes stress granule assembly to facilitate pancreatic carcinogenesis

**DOI:** 10.1101/2025.09.25.678687

**Authors:** Xue Luan, Jingke Ding, Yifu Zhu, Yan Huang, Na Wang, Chong Hong, Yanli Cheng, Ronghua Yu, Meng Wang, Zhongjun Zhao, Liye Ma, Jing Xue, Wenhui Zhang, Jintao Xu, Quanda Li, Yuran Zhao, Huixia Zhang, Kunkun Tu, Yong Liu, Jin Bai, Wenjie Ge

**Affiliations:** Cancer Institute, Xuzhou Medical University, 209 Tongshan Road, Xuzhou, Jiangsu, 221004, China; Center of Clinical Oncology, The Affiliated Hospital of Xuzhou Medical University, 99 West Huaihai Road, Xuzhou, Jiangsu, 221002, China; Jiangsu Center for the Collaboration and Innovation of Cancer Biotherapy, Xuzhou Medical University, 209 Tongshan Road, Xuzhou, Jiangsu, 221004, China; School of Life Science, Shanxi Normal University, Taiyuan, Shanxi, 030000, China; Jiangsu Key Laboratory of Coal-based Greenhouse Gas Control and Utilization, China University of Mining and Technology, Xuzhou, Jiangsu, 221008, China; Carbon Neutrality Institute, China University of Mining and Technology, Xuzhou, Jiangsu, 221008, China

## Abstract

G3BP1 promotes pancreatic ductal adenocarcinoma (PDAC) tumorigenesis driven by oncogenic *KRAS* mutations through liquid-liquid phase separation (LLPS)-mediated assembly of stress granules (SGs). However, the regulatory mechanisms remain elusive. We identify the antioxidant enzyme NAD(P)H quinone dehydrogenase 1 (NQO1) as a novel SG regulator that enhances SG assembly in pancreatic cancer cells independently of reactive oxygen species (ROS). Mechanistic studies reveal that NQO1 does not regulate the expression of *G3BP1* or other SG-associated genes. Instead, NQO1 undergoes LLPS dependent on its RNA-binding K homology (KH) like domain. Further analysis demonstrates that residues 121–131 of NQO1 directly interact with G3BP1’s RNA-binding domain (RBD), enhancing the multivalency of G3BP1 complexes to potentiate LLPS-driven SG assembly. This interaction accelerates cell proliferation, KRAS mutation-induced acinar-to-ductal metaplasia (ADM), and pancreatic carcinogenesis. Notably, the interaction between NQO1 residues 121–131 and G3BP1’s RBD is essential for NQO1 phase condensation under stress condition. Integrative analysis of human PDAC transcriptomic datasets reveals a weak association between *NQO1* and *G3BP1* levels. Importantly, both NQO1 and G3BP1 formed biological condensates and co-localized in the lesions of human PDAC tissue sections. Our study uncovers a novel *KRAS* mutation-driven mechanism of pancreatic carcinogenesis through the lens of phase separation, transcending conventional gene expression regulation and offering new insights into non-canonical KRAS-NQO1-G3BP1-SG regulatory networks in PDAC initiation.

## Introduction

Oxidative stress, characterized by the excessive accumulation of ROS^1^, plays a dual role in maintaining cellular homeostasis and promoting tumor initiation^2, 3^. Notably, Nuclear Factor Erythroid 2-Related Factor 2 (Nrf2), the master oxidative stress regulator, is dysregulated and promotes carcinogenesis and progression in pancreatic cancer^4–6^. In our previous study, we elucidated the molecular mechanisms underlying Nrf2 dysregulation in cancer cells^7^. NQO1, the predominant antioxidant target gene of Nrf2^8, 9^, its expression levels gradually accumulated during the initiation and development of PDAC^10^ and high expression of NQO1 predicted to reduce survival time of PDAC patients^11^. However, the precise biological significance of NQO1 in PDAC remains poorly understood.

Oncogenic *KRAS* mutation serves as the master driver of pancreatic carcinogenesis^12–14^. Recent studies demonstrate that mutant KRAS induces abnormal SG aggregation in ADM and pancreatic intraepithelial neoplasia (PanIN) lesions in both murine models and human PDAC tissues^15^, suggesting SG involvement in PDAC initiation. G3BP1/SG complexes further promote malignant progression of PDAC^16^, and pharmacological targeting of SGs has been shown to significantly inhibit pancreatic carcinogenesis in murine models^17^. However, the precise mechanisms by which mutant KRAS drives SG assembly and regulates PDAC carcinogenesis remain elusive.

The assembly of molecular condensates in cells, including SGs, is predominantly mediated through LLPS. This process regulates key cellular processes such as RNA metabolism, gene expression, and signal transduction pathways^18–21^. G3BP1 acts as a molecular switch for SG assembly by recruiting RNA and RNA-binding proteins (RBPs) to establish multivalent intermolecular interactions, which is essential for phase separation^22–24^. However, the precise regulatory mechanisms governing G3BP1-dependent stress granule assembly remain incompletely characterized.

In this study, we demonstrate that NQO1 is a novel regulator of G3BP1/SG dynamics, promoting SG assembly in a ROS-independent manner in PDAC cells. Although NQO1 does not modulate the expression of *G3BP1* and other SG-associated gene, it enhances G3BP1-driven LLPS via protein-protein interactions. NQO1 has been reported as a potential RBP^25^, and we found NQO1 contains one RNA binding KH like domain, which is critical for NQO1 LLPS. The interaction between NQO1 (121-231 residues) and of G3BP1 (RBD domain) enhances the multi-valency of G3BP1 condensates, thereby promoting G3BP1-mediated LLPS and SG assembly. Interestingly, the interaction between NQO1 and G3BP1 is essential for NQO1 phase condensation under stress conditions. Importantly, the *Kras^G12D^*mutation promotes *NQO1* expression, and genetic ablation of *nqo1* significantly attenuates pancreatic carcinogenesis and suppresses SG formation in *Kras^G12D^, Pdx1^Cre^*(KC) mice and 3D acinar organoids. We further observed a weak association between *NQO1* and *G3BP1* expression profiles in human PDAC samples. Both NQO1 and G3BP1 were phase separated and co-localized in the lesions of human PDAC samples. Collectively, our findings elucidate the molecular mechanisms by which *KRAS* mutations drive SG biogenesis and pancreatic carcinogenesis, providing a novel theoretical framework for early diagnosis and therapeutic intervention of PDAC.

## Results

### NQO1 is a novel SG regulator, promoting SG assembly in PDAC cells independent of ROS

To determine the biological function of NQO1 in PDAC, we used Mia-PaCa2 cells, which have a relatively high expression of NQO1 (Extended Data Fig. 1a), as the research subjects. Transfected with Flag-tagged NQO1 followed by affinity purification-coupled liquid chromatography-mass spectrometry (LC-MS/MS) analysis (Fig. 1a). Proteomic profiling identified 202 interacting proteins. Gene Ontology (GO) enrichment analysis revealed significant association with mRNA processing (GO:0006397), mRNA splicing, via spliceosome (GO:0000398), and SG assembly (GO:0034063) (Fig. 1b). SG, membrane-less ribonucleoprotein condensates, are increasingly recognized as critical mediators of cellular adaptation to stress through selective sequestration of untranslated mRNAs and signaling molecules (e.g., TIA-1, G3BP1)^26, 27^. Venn analysis demonstrated 57 overlapping proteins between NQO1 interactors and known SG constituents^22^ (Fig. 1c), with GO terms predominantly associated with SG assembly (GO:0034063) (Fig. 1d). The enriched SG assembly (GO:0034063) includes protein such as G3BP1, G3BP2, and TIA1 et.al (Fig. 1e). Given the diversity of RNA granules including P-bodies and SG^28, 29^, we examined NQO1 co-localization with G3BP1/SG under various stressors including heat shock (42°C, 2h), thapsigargin (Tg, 30 μM, 6h), and sodium arsenite (NaAsO_2_, AS, 1 mM, 1h) treatment in Mia-PaCa2 cells. Immunofluorescence confirmed significant NQO1-G3BP1 co-localization across all stress conditions, whereas minimal co-localization occurred with P-body marker DCP1A (Extended Data Fig. 1b), suggesting NQO1 may specifically regulate SG dynamics (Fig. 1f).

**Fig. 1.**
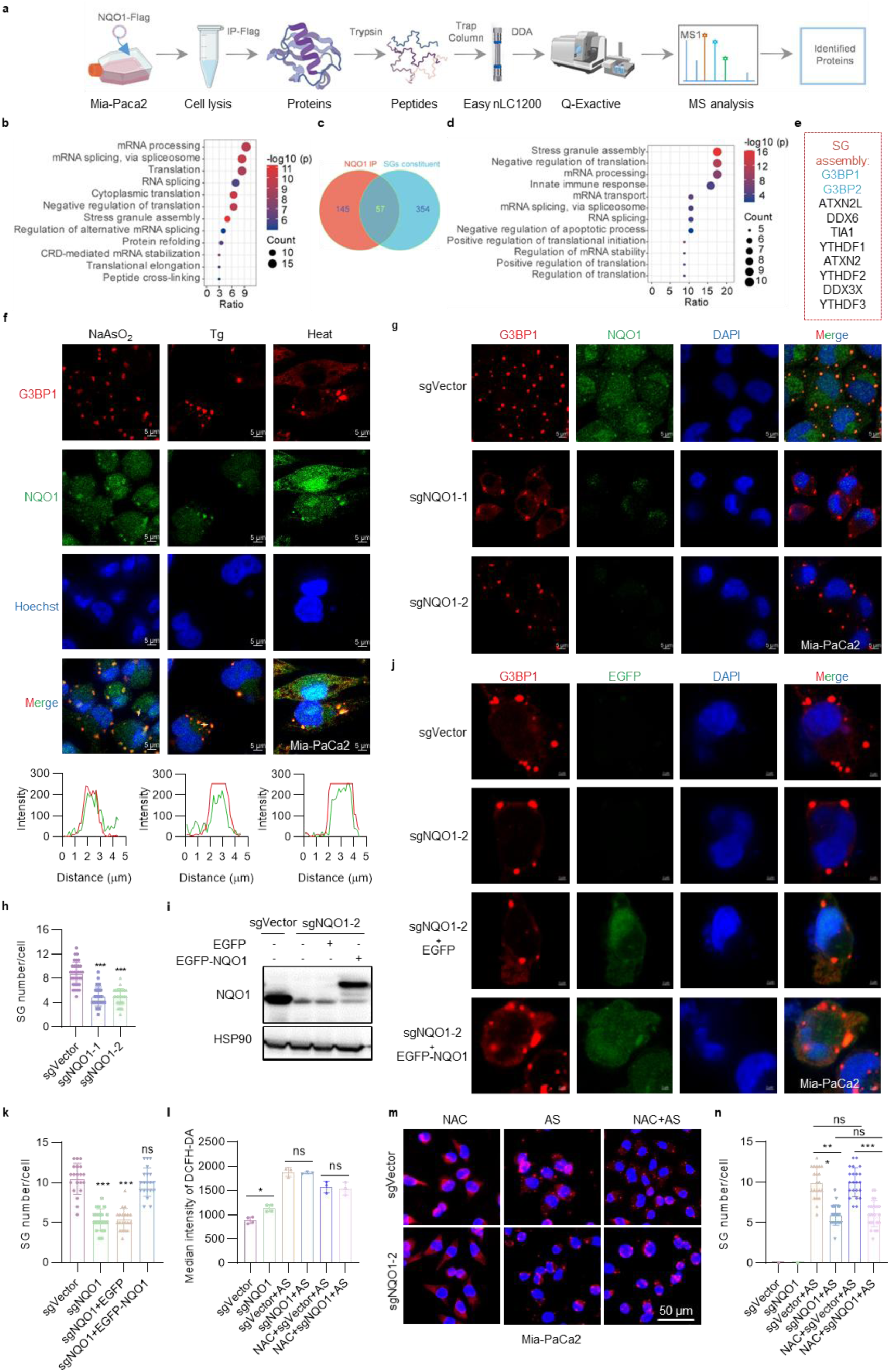
NQO1 is a novel SG regulator, promoting SG assembly in PDAC cells independent of ROS. **a,** Schematic representation of the interactive protein with NQO1 screening strategy. **b,** Functional enrichment analysis of NQO1 interacting protein is performed by using DAVID. **c,** Venn analysis of the interacting proteins and proteins related to SG regulation. **d,** Functional categorization of the 57 genes in Fig. 1c through GO enrichment analysis. **e,** Depiction of protein interactome mediating NQO1-dependent stress granule biogenesis. **f**, Representative confocal images of NQO1 clusters and SG visualized by IF staining of NQO1 (green) along with G3BP1 (red) in Mia-PaCa2 cells treated with NaAsO_2_ at 1 mM for 1 h, Tg at 30 μM for 6 h or 42 °C for 2 h. Nucleus was stained with Hoechst (blue). Scale bar, 5 μm. White dashed lines indicate the co-localization and the fluorescence intensities is shown on the underneath. Results are presentative of three independent experiments for each condition. **g-h,** Vector, *NQO1* KO Mia-PaCa2 cells were treated with NaAsO_2_ (1 mM, 1 h) and immune fluorescence stained for G3BP1 to quantify SG. SG numbers were calculated from three independent replicates of at least 20 cells, and *p* value were determined by unpaired Student’s t test, ***, *p*<0.001. Scale bar, 10 μm. **i,** Western blotting showing the rescued levels of *NQO1* in *NQO1* KO Mia-PaCa2 cells. HSP90 used as the protein loading control. **j-k,** *NQO1* KO Mia-PaCa2 cells were infected with pcSslenti-EF1-EGFP-P2A-Puro-CMV-MCS-3×FLAG-WPRE or pcSslenti-EF1-EGFP-P2A-Puro-CMV-NQO1-3 × FLAG-WPRE lentivirus plasmid and immune stained with G3BP1 to quantify SG. SG numbers were calculated from three independent replicates of at least 20 cells, *p* value was determined by unpaired Student’s t test. ***, *p* < 0.001, ns, not significant, *p* >0.05. Scale bar, 2 μm. **l,** DCFH-DA was used to detect ROS in Vector and *NQO1* KO Mia-PaCa2 cells after NaAsO_2_ (1 mM, 1 h) treatment alone or with NAC (1 mM, 12 h) (n=3). Error bars represent mean ± s.d. Data analyzed by unpaired Student’s t test. *, *p* < 0.05, ns, not significant, *p* >0.05. **m,** Immunofluorescence staining of G3BP1 was performed to assess the effect of *NQO1* ablation and NAC treatment on SG assembly under normal and stress. Scale bar, 50 μm. **n,** SG numbers were calculated from three independent replicates of at least 20 cells, and *p* value were determined by unpaired Student’s t test.

To validate NQO1’s role in SG assembly, CRISPR-Cas9-generated *NQO1* knockout (KO) Mia-Paca2 and CFPAC-1 cells (Extended Data Fig. 1c-d) were subjected to AS treatment. SG quantification by immunofluorescence revealed around 50% reduction of SG formation in *NQO1*-KO cells (Fig. 1g-h, Extended Data Fig. 1e-f). Overexpression of EGFP-NQO1 in *NQO1* KO Mia-PaCa2 cells almost completely restored SG capacity (Fig. 1i-k), confirming NQO1 as an important novel SG regulator in PDAC cells.

Considering the predominant ROS-dependent oncogenic role of NQO1 in cancer cells^8, 30, 31^, we assessed whether its regulatory effect on SG formation is mediated through ROS. The classical antioxidant N-acetylcysteine (NAC) was employed to neutralize intracellular ROS, with DCFH-DA staining used to measure ROS levels and immunofluorescence to monitor SG formation. Genetic ablation of *NQO1* elevated basal ROS in Mia-PaCa2 cells, while no significant ROS enrichment was observed in *NQO1*-KO cells upon AS treatment. ROS levels were attenuated by 1-hour pretreatment with 5 mM NAC (Fig. 1l). However, ROS scavenging by NAC did not alleviate the NQO1 deficiency-induced SG formation induced by AS (Fig. 1 m,n). Collectively, our data indicate that NQO1 promotes SG formation in PDAC cells through ROS-independent mechanism.

### NQO1 undergoes LLPS

To investigate the molecular basis of NQO1-mediated SG biogenesis in PDAC cells, we conducted RNA sequencing on *NQO1* KO and related control Mia-PaCa2 cells exposed to 1 mM AS for 1 h. Differential gene expression analysis was conducted and identified 1837 differentially expressed genes (DEGs; adjusted *p* < 0.05, fold change > 2) between *NQO1* KO and control groups. Intersectional analysis of these DEGs with established SG core components^22^ and NQO1 RNA immunoprecipitation (RIP)-enriched genes^25^ demonstrated no overlap (Fig. 2a). Subsequent protein level validation in Mia-PaCa2 and CFPAC-1 cells revealed no discernible alterations in G3BP1 expression at either mRNA (qRT-PCR, *p* ≥ 0.73) or protein levels (Extended Data Fig. 2a-c) following *NQO1* ablation (Fig. 2b). Indicating that NQO1 regulates SG assembly independent of transcriptional control over core SG machinery.

**Fig. 2.**
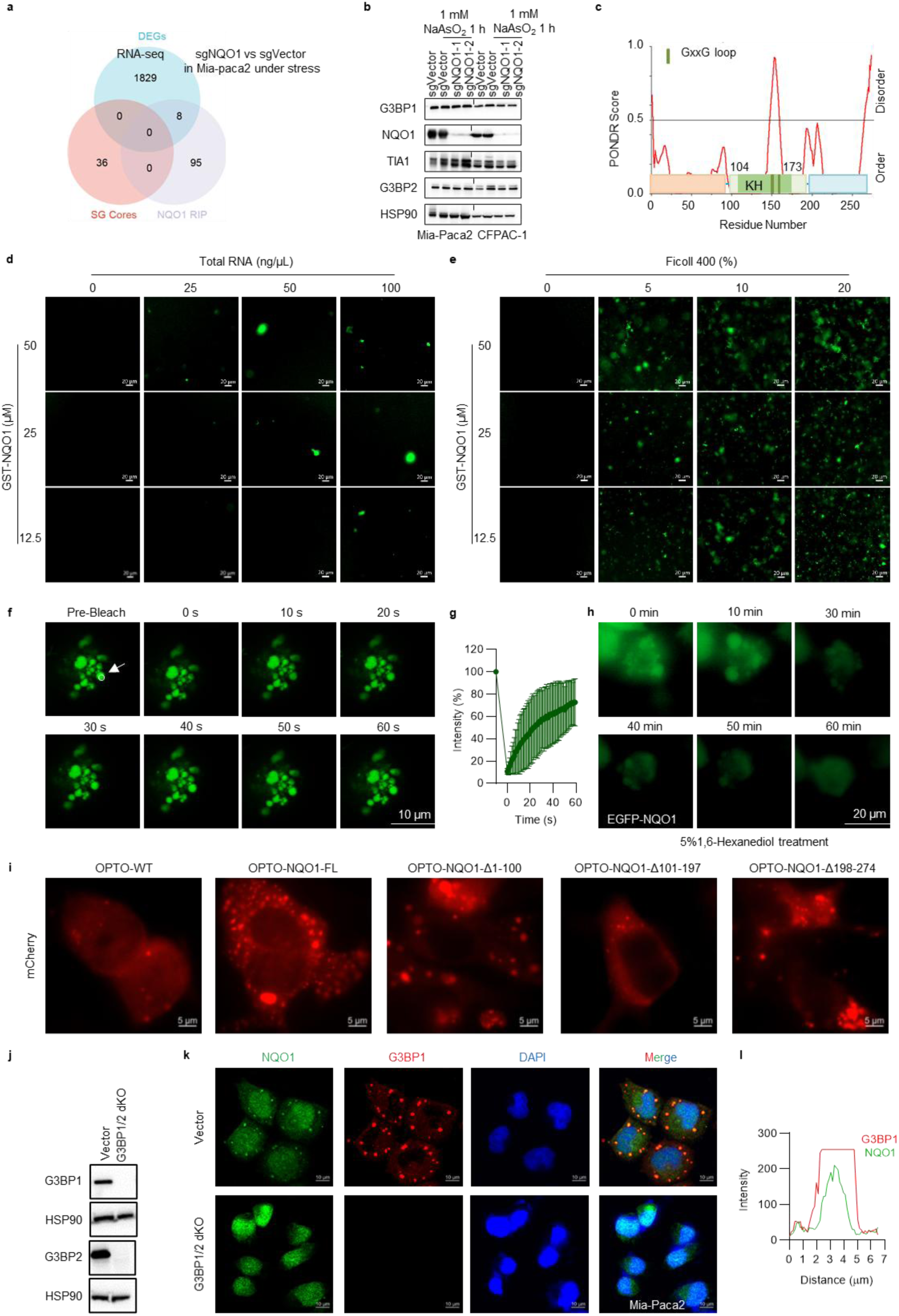
NQO1 undergoes LLPS. **a,** RNA-sequencing of *NQO1* KO and related control Mia-PaCa2 cells exposed to 1 mM AS (1 h) revealed differentially expressed genes (DEGs). Comparative analysis using Venn diagrams showed no intersection between these genes, SG core components, and NQO1 RIP-seq targets. **b,** Western blotting was performed to examine G3BP1, G3BP2, and TIA1 protein expression in *NQO1* KO Mia-PaCa2 and CFPAC-1 cells following 1 mM NaAsO_2_ treatment (1 h). HSP90 was the protein loading control. **c,** Intrinsic disorder prediction using PONDR Score identified three distinct regions in NQO1 protein (scores >0.5 indicate disordered characteristics). The KH domain is consist with 104-173 residues. **d,** Representative fluorescence images show 488-NHS ester-labeled GST-NQO1 protein droplets formed at varying concentrations in the present of indicated total RNA solution. Scale bar 20 µm. **e,** Phase-separated droplets of 488-NHS-labeled GST-NQO1 were imaged in Ficoll 400-based crowding environments, with concentration-dependent effects documented through representative microscopy. Scale bar 20 µm. **f,** Mia-PaCa2 cells transfected with pcSlenti-CMV-NQO1-linker-EGFP-3×Flag-PGK-WPRE3 plasmid were subjected to fluorescence recovery after photo-bleaching (FRAP) using a Leica STELLARIS 5 confocal microscope (488 nm channel). Time-lapse imaging captured EGFP-NQO1 droplet recovery under 1 mM NaAsO_2_ treatment. Scale bar 10 µm. **g,** Triplicate experiments quantified fluorescence intensity changes over time. **h,** Time-lapse recordings from live-cell stations captured 1,6-hexanediol (5%)-mediated dispersal of EGFP-NQO1 condensates under 1 mM NaAsO_2_ stress in lentivirus-transduced Mia-PaCa2 cells. Scale bar 20 µm. **i,** HEK-293T cells expressing pHR-mCherry-Cry2-NQO1 fusion constructs and truncation mutants were photo-activated with 488 nm light (1 s pulse at minimal intensity). mCherry fluorescence was monitored post-activation. Scale bar 5 µm. **j,** Western blot analysis confirmed complete ablation of *G3BP1* and *G3BP2* protein expression in the double knockout (dKO) Mia-PaCa2 cells. HSP90 was the protein loading control. **k,** Representative immunofluorescence images show NQO1 puncta formation in vector control versus *G3BP1/2* dKO Mia-PaCa2 cells following 1 mM NaAsO_2_ treatment (1 h). Scale bar 10 μm. **l,** White dashed lines in panel k denote regions of G3BP1-NQO1 co-localization intensity analysis.

To determine the mechanism of NQO1-SG regulation, we analyzed the NQO1 protein structure based on the *in-silicon* PONDR prediction method. We found that NQO1 contains an IDR region featured with two conserved GXXG loops (150-153 aa,160-163 aa) (Fig. 2c). These GXXG loops are characteristic of the classical RNA-binding KH domain^32, 33^. Complementary to existing evidence positioning NQO1 had RNA binding activity^25^, we identified that the 104-173 amino acid segment of NQO1 adopts a β-α-α-α-β-α topology, which displays a single β-strand replacement by an α-helix compared to the canonical type II KH domain (β-α-α-β-β-α). These observations suggest that NQO1 may share KH-like domain functionality. NQO1 contains one KH like domain (104-173 aa) further supports the notion that NQO1 is an RNA binding protein. Since most RNA binding proteins were intrinsically disordered^34^, then we postulated that this structural configuration might enable phase-separation competence. Experimental validation commenced with prokaryotic expression and purification of GST-tagged NQO1, followed by covalent labeling using 488-NHS ester fluorophores to enable phase-separation tracking. Remarkably, GST-NQO1 exhibited robust droplet formation in vitro when exposed to RNA and crowding agents (Ficoll 400), as evidenced by fluorescent condensates (Fig. 2d,e, Extended Data Fig. 2d,e). Cellular overexpression studies in AS-treated Mia-PaCa2 cells showed EGFP-NQO1 foci with rapid fluorescence recovery post-bleaching (Fig. 2f,g), characteristic of liquid-like phase-separated condensates. Treatment with 1,6-hexanediol caused time-dependent dissolution of EGFP-NQO1 puncta (Fig. 2h), further supporting LLPS activity. Collectively, these data demonstrate that NQO1 undergoes LLPS both *in vitro* and *in vivo*.

Our investigation into LLPS-critical residues of NQO1 using PSPHunter identified three candidate regions (46-72, 77-97, and 121-154) through computational prediction^35^ (Extended Data Fig. 2f). Guided by secondary structure preservation principles, truncation sites were strategically designed at residues 100 and 197 to avoid disrupting β-sheet/α-helix configurations (Extended Data Fig. 2f). Using the optogenetic vector pHR-mCh-Cry2WT (Opto-WT, Addgene #101221) as template, we generated Opto-NQO1-FL through Gibson assembly-based recombination. Sequential domain deletions yielded three constructs: Opto-NQO1-Δ1-100 (N-terminal truncation), Δ101-197 (central KH like domain removal), and Δ198-274 (C-terminal truncation). Transfected HEK-293T cells underwent 488 nm blue light stimulation (1s duration) for optogenetic induction, with phase-separated condensates captured via mCherry fluorescence microscopy. These results revealed thatΔ101-197 caused significant attenuation of LLPS capacity (Fig. 2i). These findings indicating that the 101-197 residue was required for NQO1 phase separation. Physiological NQO1 exhibits diffuse distribution but coalesces into SG. Co-localization analysis under G3BP-proficient and -deficient conditions revealed that NQO1 puncta formation during stress requires G3BP scaffold (Fig. 2j-l). Collectively, we found that the KH like domain and G3BP are critical for NQO1 LLPS.

### NQO1 interacts with G3BP1, thereby promoting G3BP1 LLPS and SG formation in PDAC cells

While SG formation is known to correlate with expression levels of core scaffold proteins^16, 20, 29^, we observed that NQO1 does not regulate canonical SG components such as G3BP1, G3BP2 and TIA1 (Fig. 2b). Given that LLPS of scaffolding proteins constitutes a fundamental mechanism for SG assembly enabling spatial organization of RNA-binding protein and RNA molecules into microscopically detectable condensates^22^. We investigated the potential regulatory role of NQO1 in this process. LC-MS/MS proteomics identified a potential novel NQO1-G3BP1 interaction, prompting us to hypothesize that NQO1 enhances G3BP1-mediated LLPS through direct binding. To validate this hypothesis, we first confirmed the direct interaction between NQO1 and G3BP1 using co-immunoprecipitation (Co-IP) and GST pull-down assays (Fig. 3a-c). To examine the regulatory role of NQO1 in G3BP1-mediated LLPS, we initiated experiments using NHS-488 ester-labeled His-G3BP1 to assess phase-separated droplet formation. Consistent with prior findings, G3BP1 exhibited LLPS under both RNA-enriched and Ficoll 400-simulated crowded environments (Fig. 3d, Extended Data Fig. 3a-c). We established the threshold LLPS concentration of G3BP1 as 25 μM with 25 ng/μL total RNA, which reduced to 12.5 μM when 5% Ficoll 400 was applied to mimic intracellular crowding. Co-incubation with 25 μM GST-NQO1 (non-phase-separating at this concentration, Fig. 2d,e) significantly amplified G3BP1 LLPS activity, with more pronounced effects observed in RNA-supplemented and Ficoll 400-crowded systems (Fig. 3e,f). Parallel controls with GST protein at matched concentrations showed negligible effects on G3BP1 phase separation (Extended Data Fig. 3d,e). These *in vitro* reconstitution experiments demonstrate that NQO1 enhances G3BP1’s phase-separation capacity at critical thresholds, thereby establishing that NQO1 promotes SG assembly in PDAC cells via augmentation of core scaffolding protein LLPS initiation.

**Fig. 3.**
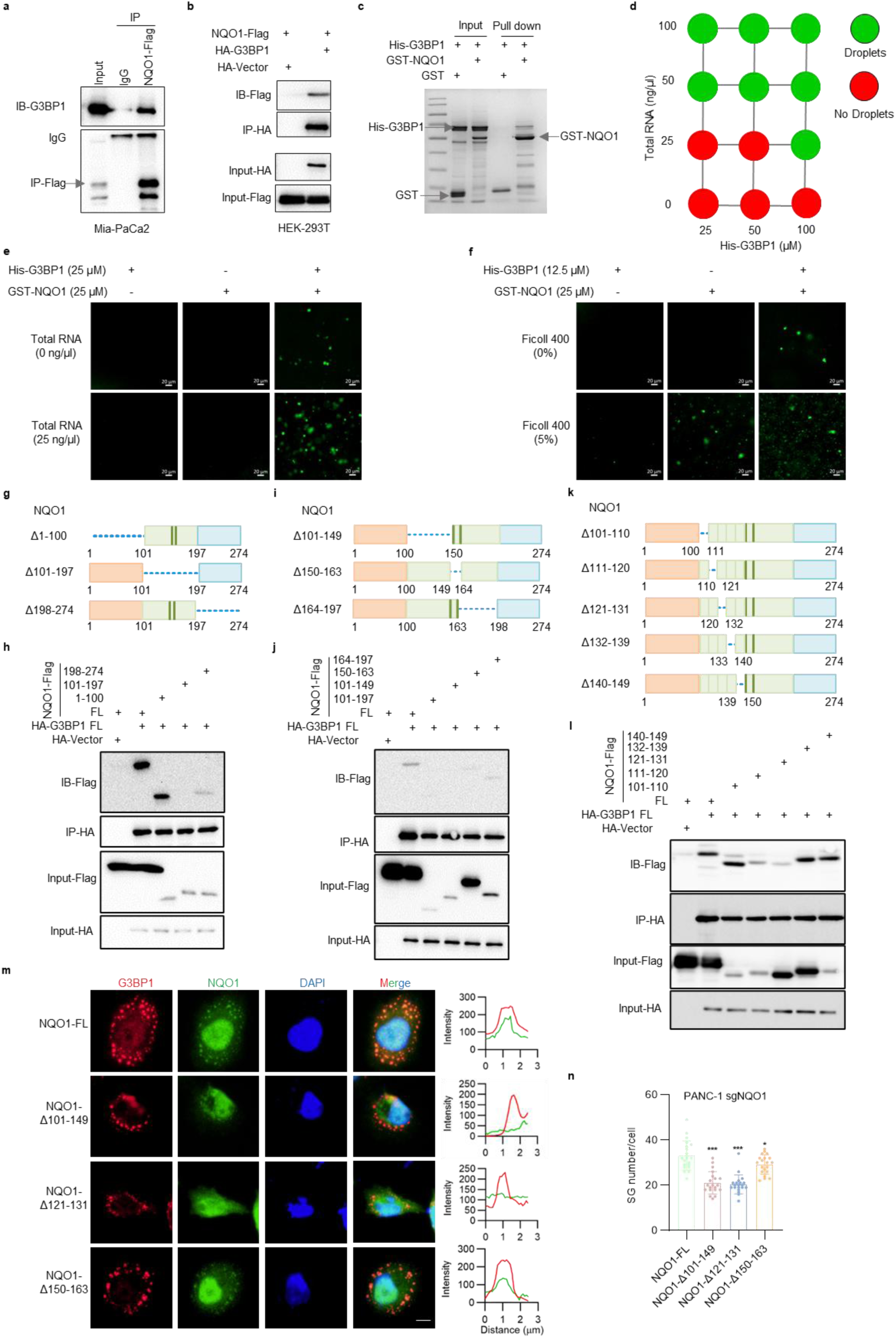
NQO1 interacts with G3BP1, thereby promoting G3BP1 LLPS and SG formation in PDAC cells. **a,** Mia-PaCa2 cells were transfected with pCDNA3.1-NQO1-Flag plasmid for 48 h. Cell lysates were subjected to co-immunoprecipitation using Protein A/G agarose beads conjugated with Flag antibody, with IgG antibody as negative control. Western blotting was performed to analyze the interaction between Flag-NQO1 and endogenous G3BP1. **b,** Lysates from PKH3-HA and pCDNA3.1-NQO1-Flag or PKH3-G3BP1-HA and pCDNA3.1-NQO1-Flag co-transfected HEK-293T cells underwent magnetic bead-based co-IP with Flag antibody. G3BP1 binding to Flag-NQO1 was validated by immunoblotting. **c,** Purified GST or GST-NQO1 bait proteins were incubated with GSH-agarose beads at 4°C for 2 h. His-G3BP1 prey protein was then added for overnight incubation. After extensive washing, elution with GSH-containing buffer was performed. SDS-PAGE assessed direct interaction between GST-NQO1 and His-G3BP1. **d,** Summary of G3BP1 phase separation behaviors in the presence of total RNA. Representative images are shown in Extended Data Fig. 3a. **e-f.** Representative images of NHS-488 ester-labeled full-length G3BP1 phase separation at critical concentration with GST-NQO1 addition. Conditions with/without total RNA (e), with/without Ficoll 400 (f). Scale bar 20 μm. **g-h,** Co-IP using HA-conjugated magnetic beads was performed on lysates from HEK-293T cells co-expressing HA-G3BP1 with various NQO1 truncations (pCDNA3.1-NQO1-3×Flag, pCDNA3.1-NQO1Δ1-100-3×Flag, pCDNA3.1-NQO1Δ101-197-3×Flag, pCDNA3.1-NQO1Δ198-274-3×Flag). Immunoblotting revealed differential binding patterns. **i-j,** HEK-293T cells co-expressing HA-G3BP1 FL with NQO1 FL/Δ101-197/Δ101-149/Δ150-163/Δ164-197-Flag plasmids underwent HA bead-based co-IP. Western blotting determined interaction between NQO1 subdomains and G3BP1. **k-l,** Co-transfection of PKH3-HA-G3BP1 FL with pCDNA3.1-NQO1 FL/Δ101-110/Δ111-120/Δ121-131/Δ132-139/Δ140-149-Flag plasmids in HEK-293T cells was followed by HA beads co-IP. Immunoblotting delineated critical residues of NQO1 required for G3BP1 binding. **m,** *NQO1* KO PANC-1 cells transfected with NQO1 FL/truncation mutants for 48 h, followed by treatment with 1 mM NaAsO₂ for 1 hour. SG formation was quantified by immunofluorescence. Scale bar 10 μm. **n,** Quantitative analysis of SG numbers (≥20 cells/group) demonstrated differential responses among NQO1 variants (unpaired Student’s t test, *, *p* < 0.05, ***, *p* < 0.001).

To systematically investigate the key motif of NQO1-mediated SG regulation, we generated a series of Flag-tagged NQO1 mutants through truncation mutagenesis (Fig. 3g, h). Co-immunoprecipitation mapping revealed that the 101-197 residues serve as the critical binding module for G3BP1 interaction (Fig. 3h). Subsequent segmentation based on GXXG loops topology further localized the functional interface to residues 101-149 (Fig. 3 i,j). Through five rationally designed truncations preserving secondary structure integrity (Fig. 3k), comprehensive binding analysis identified residues 121-131 as the minimal core motif essential for G3BP1 protein interaction (Fig. 3l).

Our experimental data demonstrate that NQO1 is G3BP-dependent during AS-induced LLPS, as shown in Figure 2k. Based on this, we propose that the structural domain mediating the NQO1-G3BP1 interaction is a critical determinant of stress-induced NQO1 LLPS formation. Domain-specific rescue experiments conducted in *NQO1* KO PANC-1 cells (Extended Data Fig. 3f) compared four constructs: NQO1 FL, Δ101-149 (disrupting binding interface), Δ121-131 (core interaction domain deletion), and Δ150-163 (GXXG loop truncation affecting RNA binding). Notably, the Δ101-149 and Δ121-131 truncations completely abolished NQO1-G3BP1 co-localization, resulting in complete NQO1 LLPS failure, compared to FL controls (Fig. 3m). Although Δ150-163 retained the capacity for NQO1-G3BP1 co-localization (Fig. 3m), the 12% decrease in SG formation (p<0.05 vs FL; Fig. 3n) suggests that RNA-binding activity may play a non-essential but supportive role in G3BP1 LLPS potentiation.

These findings indicate that residues 121-131 of NQO1 constitute an essential domain for both interaction with G3BP1 and autonomous liquid-liquid phase separation (LLPS). This structural-functional relationship provides a molecular basis for targeting NQO1-G3BP1/ SG interactions in the context of therapeutic development.

### The RBD domain of G3BP1 was critical for NQO1 interaction

To systematically characterize G3BP1-NQO1 binding domain, we generated HA-tagged G3BP1 truncation mutants (Fig. 4 a,c) and performed co-IP assays. Notably, deletion of G3BP1’s RBD (RGG and RRM) domain abolished its interaction with NQO1 (Fig. 4 b,d, Extended Data Fig. 4a). While EGFP-RGG fusion constructs confirmed direct binding to the RGG motif (Fig. 4e). Evolutionary conservation of this interaction was evidenced by NQO1’s capacity to bind with RGG-containing proteins Caprin1 and UBAP2L (Fig. 4f). To confirm the interaction specificity between NQO1 and RRM domain, we performed binding assays with TIA1 (exclusively RRM domain) and FUS protein (containing both RRM and RGG domains). Results demonstrated that NQO1 binds to FUS protein with both RGG and RRM motifs, as well as TIA1 protein that solely retains the RRM domain. (Fig. 4g). These results suggest that the universal role of NQO1 interacting with RBD domain of G3BP1.

**Fig. 4.**
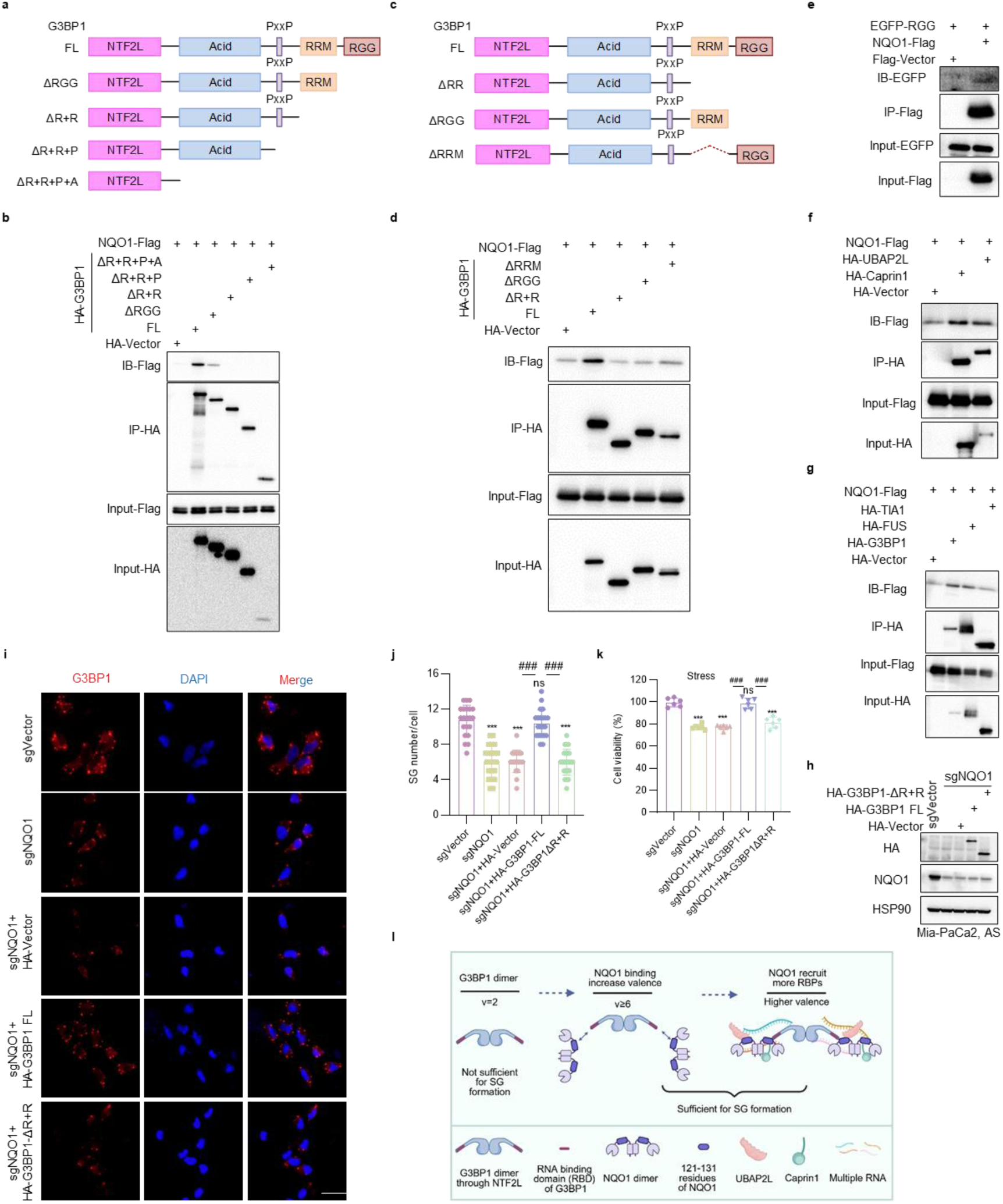
The RBD domain of G3BP1 was critical for NQO1 interaction. **a,** The cloning strategy HA-G3BP1 truncation mutants were generated. **b,** pCDNA3.1-NQO1-Flag was co-expressed with PKH3-HA-Vector, PKH3-HA-G3BP1 FL, or PKH3-HA-G3BP1 truncation mutants in HEK-293T cells. Post 48-hour incubation, Co-IP assays using HA-tagged magnetic beads were performed on cell lysates. Western blotting was employed to delineate the interaction domains of NQO1 with G3BP1. **c,** PKH3-HA-G3BP1 FL, PKH3-HA-G3BP1-ΔRRM+RGG (HA-G3BP1-ΔR+R), PKH3-HA-G3BP1ΔRGG and PKH3-HA-G3BP1ΔRRM mutants were constructed. **d,** Co-expression of pCDNA3.1-NQO1-Flag with HA-Vector, HA-G3BP1 FL, or truncation mutants in HEK-293T cells was followed by Co-IP assays with HA antibody-coupled beads at 48 h post-transfection. Subsequent Western blotting precisely mapped the binding residues of NQO1-G3BP1 interaction. **e,** Co-IP experiments in HEK-293T cells were conducted to investigate the binding of NQO1 to the RGG domain. **f,** Co-IP experiments in HEK-293T cells were conducted to investigate the binding of pCDNA3.1-NQO1-Flag to PKH3-HA-UBAP2L and PKH3-HA-Caprin1 protein. **g,** Co-IP experiments in HEK-293T cells were conducted to investigate the binding of NQO1 to TIA1 and FUS protein. **h,** The expression levels of PKH3-HA-G3BP1 FL and PKH3-HA-G3BP1 ΔR+R in *NQO1* KO Mia-PaCa2 cells under AS treatment for 1 hour, the expression were determined by Western blot analysis. HSP90 was the protein loading control. **i,** Representative immunofluorescence images showing SG formation in *NQO1* KO Mia-PaCa2 cells overexpressing PKH3-HA-G3BP1 FL or PKH3-HA-G3BP1 ΔR+R. Scale bar 20 μm. **j,** Statistical analysis of SG counts in Fig. 4i, representing three biological replicates with a minimum of 20 cells analyzed per condition. **k,** Cell proliferation assays of *NQO1* KO Mia-Paca2 cells transfected with PKH3-HA-Vector, PKH3-HA-G3BP1 FL, or PKH3-HA-G3BP1 ΔR+R under AS-induced conditions (n=6 independent experiments). Data were analyzed using unpaired Student’s t test. ***, *p* < 0.001; ns, not significant, *p* > 0.05; compared to sgVector; ^###^, *p* < 0.001; relative to sgNQO1+HA-G3BP1 FL group. **l,** The schematic model depicting NQO1 enhanced G3BP1 condensate valence to promote its LLPS. The dimeric G3BP1 (Valence = 2) is insufficient to form SG. The dimeric NQO1 binds to the RBD of G3BP1, thereby recruiting more RNA binding proteins (RBPs) and significantly increasing the complex valence, ensuring the formation of SG. Figure 3l created using BioRender.

Rescue experiments in *NQO1*-deficient Mia-PaCa2 cells demonstrated that exogenous expression of G3BP1 FL, but not the ΔR+R mutant (lacking RGG and RRM domains) (Fig. 4h), restored AS-induced SG formation (Fig. 4 i,j, n≥20 cells/group from three independent experiments). These results established that NQO1 as a critical enhancer of G3BP1 LLPS, with the RBD serving as the central regulatory node. NQO1 knockout substantially impaired Mia-PaCa2 proliferation (Extended Data Fig. 4b). Intriguingly, under basal conditions, G3BP1 FL and ΔRBD overexpression in *NQO1* KO cells (Extended Data Fig. 4c), G3BP1 FL overexpression enhanced proliferation by 13.37% compared HA-Vector control (*p* < 0.001), not fully restoring to sgVector levels (Extended Data Fig. 4d). Post-AS treatment, G3BP1 FL-expressing cells exhibited greater proliferative capacity versus HA-Vector and ΔR+R group (*p* < 0.001), achieving parity with sgVector controls (Fig. 4k). The ΔR+R mutant group showed complete loss-of-function across all experimental paradigms, with indistinguishable proliferation when compared with the HA-Vector group.

The NTF2L and RBD domains were critical for the LLPS and SG regulation of G3BP1^22–24^. Based on the patchy colloid model, the valency of a certain protein defines whether it will enhance or inhibit stress granule formation^23^. Proteins with large valencies, like UBAP2L, Caprin1, FUS bind with G3BP1 through NTF2L domain and promote the multi-valency interactions, eventually promote SG assembly. On the other hand, USP10 is a low valency protein and act as a cap to prevent SG formation by interfering G3BP1-Caprin1 interactions. The RNA binding activity of G3BP1’s RBD domain is also required for SG assembly since RNA serve as the substrate to enhance the valencies of these RNA binding proteins^23^. In this study, we deciphered that RBD not only binds with RNA but also interact with NQO1. This binding between NQO1 and RBD may blocked the G3BP1 RNA binding, while the KH domain of NQO1 will probably compensate the RNA binding capacity of the SG complex. In the meantime, the SG regulators UBAP2L, caprin1, FUS et al binds not only G3BP1 but also NQO1. Taken together, the NQO1-G3BP1-RBPs and RNA hetero-organizations increase the valencies of the SG complex, promote LLPS, facilitate the SG assembly and sustaining PDAC cell proliferation (Fig. 4l).

### Genetic ablation of *nqo1* in KC mice attenuated PDAC initiation through NQO1-G3BP1/SG axis

Our cellular-level investigations demonstrate that NQO1 facilitates malignant proliferation in PDAC cells through G3BP1/SG assembly, yet its systemic *in-vivo* biological functions remain uncharacterized. Analysis of the GSE132326 dataset revealed stage-dependent upregulation of *NQO1* during mouse PDAC progression (Fig. 5a). Western blot analysis showed negligible NQO1 expression in wild-type mouse pancreas (Extended Data Fig. 5b), suggesting potential oncogenic involvement. Single-cell transcriptomic profiling of human PDAC versus normal pancreas identified *NQO1* enrichment in ductal epithelial and vascular endothelial compartments (Extended Data Fig. 5a). To establish genetic models, we employed *Nqo1^-/-^* mice to cross with KC mice which exhibited delayed tumorigenesis (prolonged latency) and developed preneoplastic lesions histologically analogous to human PDAC precursors (Fig. 5b,c). The genotyping and expression validation confirmed successful *Nqo1* ablation in mice pancreas (Extended Data Fig. 5c; Fig. 5d), with KC mice showing NQO1 elevation versus wild-type controls (Fig. 5d). Quantitative morphometric analysis revealed significantly reduced pancreas/body weight ratios in KCN^-/-^ versus KC mice (3 months: 1.20% vs 1.64%, Extended Data Fig. 5d, p < 0.01; 6 months: 1.47% vs 2.10%, p < 0.001; Fig. 5 e,f).

**Fig. 5.**
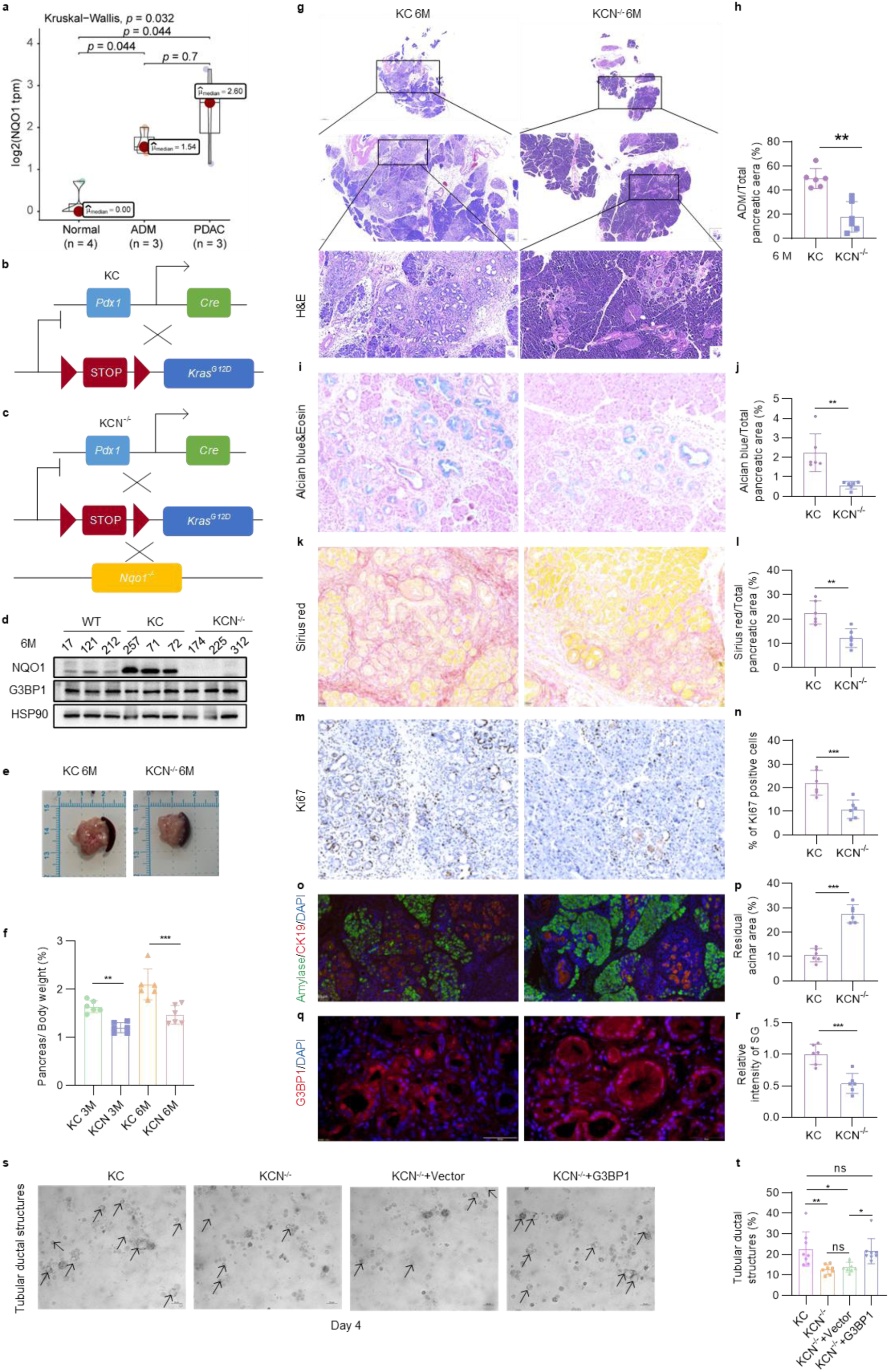
Genetic ablation of *nqo1* in KC mice attenuated PDAC initiation through NQO1-G3BP1/SG axis. **a,** Expression of *nqo1* mRNA during *Kras* mutation-driven PDAC progression (GSE132326). Inter-group differences were analyzed by Kruskal-Wallis test, with *p* < 0.05 indicating statistical significance. **b-c**, Generation strategies of KC and KCN^-/-^ mouse models. **d,** Western blot analysis of NQO1 and G3BP1 protein expression in pancreatic tissues from 6-month-old WT, KC, and KCN^-/-^ mice. HSP90 was the protein loading control. **e,** Representative pancreatic tissue morphology in 6-month-old KC and KCN^-/-^ mice. Scale bar, 1 cm. **f,** Pancreas-to-body weight ratio quantification in KC and KCN^-/-^ mice at 3 and 6 months. n=6. Mean ± s.d; **, *p* < 0.01; ***, *p* < 0.001.**g,** Hematoxylin and eosin (H&E) staining of pancreatic sections from 6-month-old KC and KCN^-/-^ mice, demonstrating 2×, 10×, and 20× magnifications (scale bars, 1000 μm, 200 μm and 100 μm respectively). **h,** Quantification of ADM as a percentage of total pancreatic area based on H&E staining. n=6. Mean ± s.d;**, *p* < 0.01. **i-j,** Alcian blue/Eosin dual staining with quantification of Alcian blue-positive areas as a percentage of total pancreatic surface. n=6. Mean ± s.d; **, *p* < 0.01. **k-l,** Sirius red staining with collagen-positive area quantification relative to total pancreatic area. n=6. Mean ± s.d; **, *p* < 0.01. **m-n,** Ki67 IHC staining with cell proliferation index calculated as Ki67-positive nuclei per total nucleated cells. n=6. Mean ± s.d; ***, *p* < 0.001. **o-p,** CK19/Amylase immunofluorescence co-staining with residual acinar area quantification (Amylase-positive regions as percentage of total pancreas). n=6. Mean ± s.d; ***, *p* < 0.001. **q-r,** G3BP1 immunofluorescence and SG enrichment analysis. Data in panels e-p were derived from pancreatic sections of 6-month-old KC and KCN^-/-^ mice (n=6). Intergroup comparisons used unpaired Student’s t-test (Mean ± s.d; ***, *p* < 0.001). **s,** Bright-field imaging of primary acinar cell clusters in collagen-based 3D culture at day 4 post-seeding. Arrows indicate ADM cells (scale bar 50 μm). **t,** Quantitative analysis of tubular ductal structures as shown in panel q (n=8, Mean ± SD; *, *p* < 0.05; **, *p* < 0.01; ns, not significant, *p* > 0.05).

Subsequent investigations focused on NQO1’s role in *Kras^G12D^*-mediated pancreatic carcinogenesis. H&E-stained sections demonstrated ADM progression occupying 4.74% and 49.75% of pancreatic area in 3-and 6-month-old KC mice, respectively, compared to 2.85% and 17.98% in age-matched KCN^-/-^ littermates (6 month: Fig. 5g,h, *p*<0.01; 3 month: Extended Data Fig. 5e,f, *p*<0.05). Consistent with attenuated ADM progression, *Nqo1* deficiency significantly reduced pathological mucin deposition (6 month: 2.24% vs KCN^-/-^ 0.57%, *p* < 0.01; Fig. 5 i,j; 3month: 0.29% vs KCN^-/-^ 0.17%, *p* > 0.05, Extended Data Fig. 5g,h) and collagen fibrosis (Sirius red quantification: 6 month:KC 22.58% vs KCN^-/-^ 12.06%, *p* < 0.01; Fig. 5k,l; 3 month: KC 3.38% vs KCN^-/-^ 2.18%, *p* > 0.05; Extended Data Fig. 5i,j). Proliferative index quantified by Ki67 immunohistochemistry (IHC) showed marked reduction in KCN^-/-^ versus KC cohorts (6 month: KCN^-/-^ 10.85% vs KC 22.04%, *p* < 0.01; Fig. 5m,n; 3month: 3.88% vs 9.37%, *p* < 0.001, Extended Data Fig. 5k,l). Amylase/CK19 dual immunofluorescence confirmed residual acinar architecture in KCN^-/-^ mice, with residual acinar areas larger than KC controls (6 month: KCN^-/-^ 27.53% vs KC 10.56%, *p* < 0.01; Fig. 5 o,p). These integrated pathophysiological analyses establish NQO1 as an important amplifier of *Kras^G12D^*-driven oncogenesis through coordinated regulation of metaplastic transformation, mucinous reprogramming, and stromal desmoplasia.

Mechanistic interrogation of the NQO1-G3BP1/SG axis revealed a 47.17% reduction in SG density in 6-month-old KCN^-/-^ mice compared to KC mice (Fig. 5 q,r, *p* < 0.001), demonstrating NQO1 dependency in maintaining SG integrity. Functional validation using 3D acinar organoid models showed significant abrogation of *Kras^G12D^*-induced ductal morphogenesis in *Nqo1^-/-^* systems (organoid formation efficiency: KC 22.67±8.37% vs KCN^-/-^ 12.70±2.32%, *p* < 0.001), which was phenotypically rescued by lentiviral G3BP1 FL overexpression (restoration to 21.61±6.13% efficiency, *p* < 0.05 vs KCN^-/-^ baseline; Extended Data Fig. 5c,m, Fig. 5s,t). These findings align with established SG assembly mechanisms, wherein G3BP1 scaffolding coordinates LLPS dynamics^20^. The integrated dataset delineates a targetable signaling nexus wherein NQO1 sustains oncogenic *Kras^mut^* signaling fidelity through mediating SG assembly to promote PDAC initiation.

### NQO1 and G3BP1 analysis in Human PDAC

Our previous findings showed that genetic ablation of *NQO1* didn’t affect G3BP1 expression but drastically repressed SG activity in both human PDAC cells and genetic modified mice (Fig.1g,2b,5d,q). To determine the expression pattern and spatial localization of NQO1 and G3BP1 in pancreatic cancer tissue sections, we first examined protein expression of NQO1 and G3BP1 in pancreatic tissues of KC mice at 1, 3, and 6 months of age. The results demonstrated age-dependent upregulation of NQO1, while G3BP1 expression showed no significant differences across time points (Fig. 6a,b). These findings further confirmed that NQO1 does not regulate G3BP1 expression. Notably, 6-month KC mice exhibited SG enrichment within pathological pancreatic tissues versus wild-type counterparts. Striking spatial co-localization between NQO1 and G3BP1 (Fig. 6c) further establishes NQO1 as an important modulator of SG in pancreatic cancer. To delineate the dynamics of *NQO1* expression during human PDAC pathogenesis, we performed transcriptomic interrogation of matched tumor-normal pairs from the TCGA cohort. This analysis revealed pronounced *NQO1* mRNA upregulation in PDAC lesions compared to histologically normal counterparts (Fig. 6d; *p* < 0.0001), with *G3BP1* exhibiting marginal yet statistically significant elevation (Fig. 6e; 2.94-fold vs. 58.48-fold for *NQO1*). Bivariate correlation analysis demonstrated weak association between *NQO1* and *G3BP1* expression profiles in human PDAC samples (Pearson’s r=0.26, Fig. 6f). The single-cell data of human PDAC show that NQO1 is mainly significantly enriched in ductal cells in PDAC tissues (Extended Data Fig. 5a). Ductal cells are the main cell characteristics for the occurrence of pancreatic cancer precursors, indicating that NQO1 may be an important trigger for the initiation of pancreatic cancer driven by *KRAS* mutations. Building on established mechanisms of oncogenic *Kras* mutation driven SG biogenesis in PanIN, we employed IHC analyses to map NQO1 induction specifically to neoplastically transformed ductal epithelium (Fig. 6g). The immunofluorescence staining results of the human pancreatic cancer tissue sections showed that NQO1 and G3BP1 were significantly enriched in the lesion area and had a significant co-localization level compared with adjacent normal tissue (ANT) (Fig. 6h). These findings demonstrated that NQO1 was a stage-specific molecular driver of PDAC initiation and progression.

**Fig. 6.**
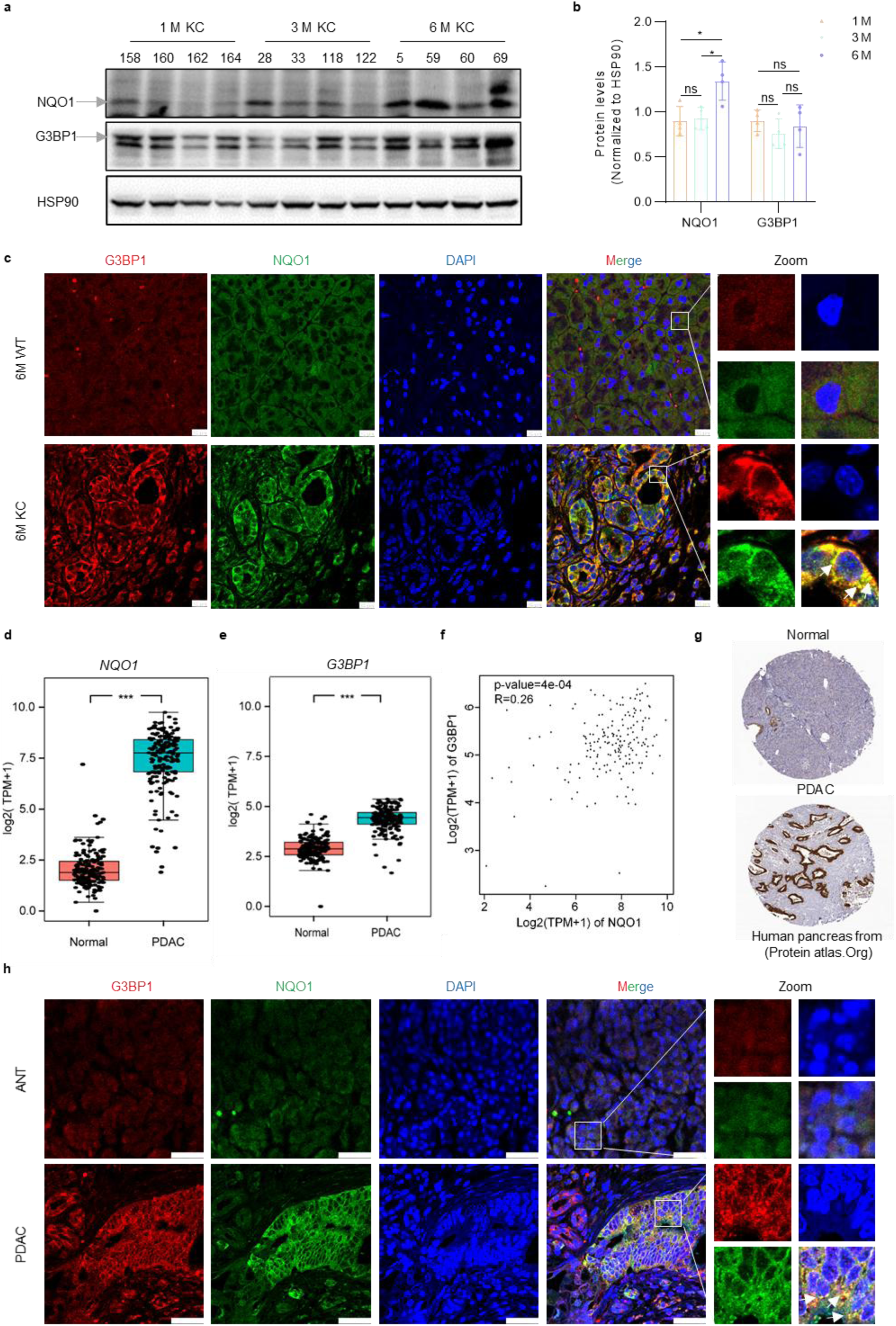
NQO1 and G3BP1 analysis in Human PDAC. **a,** Western blot analysis of pancreatic NQO1 and G3BP1 expression in KC mice at 1, 3, and 6 months post-birth. **b,** Quantitative densitometry (normalized to HSP90 loading control) of NQO1 and G3BP1 (mean ± s.d, ns, not significant, *p* > 0.05; *, *p* < 0.05, analysed by unpaired Student’s t-test) **c,** Immunofluorescence co-localization of NQO1 (green) and G3BP1 (red) in 6-month-old KC mice, with age-matched wild-type (WT) controls showing minimal overlap. Nuclei were counterstained with DAPI. Scale bar 10 μm. **d,e,** Analysis of *NQO1* and *G3BP1* mRNA expression in human PDAC samples using TCGA database. **f,** Interrogation of GEPIA co-expression network demonstrates minimal transcriptional correlation (Pearson’s R < 0.3) between *NQO1* and *G3BP1* in PDAC cohorts. **g,** IHC staining images of NQO1 in pancreatic cancer and adjacent tissues from Human Protein Atlas database. **h,** Immunofluorescence representative images for detection of co-localization levels of G3BP1 (Red) and NQO1 (Green) in human PDAC tissue and adjacent normal tissue (ANT) from human pancreatic tissue microarray. Scale bar 20 μm.

Taken together, our multimodal findings showed that oncogenic *KRAS* mutations dramatically promotes the expression of *NQO1*. In NQO1 enriched condition, NQO1 interacts with the RBD domain of G3BP1 via its 121-131 residues. Although NQO1 may occupies partial RNA-binding sites on G3BP1, its KH domain could mediate RNA interaction and the 121-131 motif recruit high valencies RBPs (e.g., UBAP2L and Caprin1, TIA1 and FUS et.al) into the SG core complex. This hetero-molecular interactions enhances the valence of SG complex, promoting G3BP1 LLPS and SG assembly. In *NQO1* deficient scenario, reduced valence of G3BP1-centered condensates impairs LLPS capacity and SG formation. *KRAS* mutation-induced SG assembly is a known driver of PDAC pathogenesis^17^. Oncogenic *KRAS* mutation promote *NQO1* expression levels, enhanced G3BP1 LLPS, SG formation, proliferation, transformation and pancreatic carcinogenesis. Collectively, the KRAS-NQO1-G3BP1/SG axis constitutes a critical regulatory network driving pancreatic carcinogenesis and may provide a novel avenue for PDAC intervention (Fig 7).

**Fig. 7.**
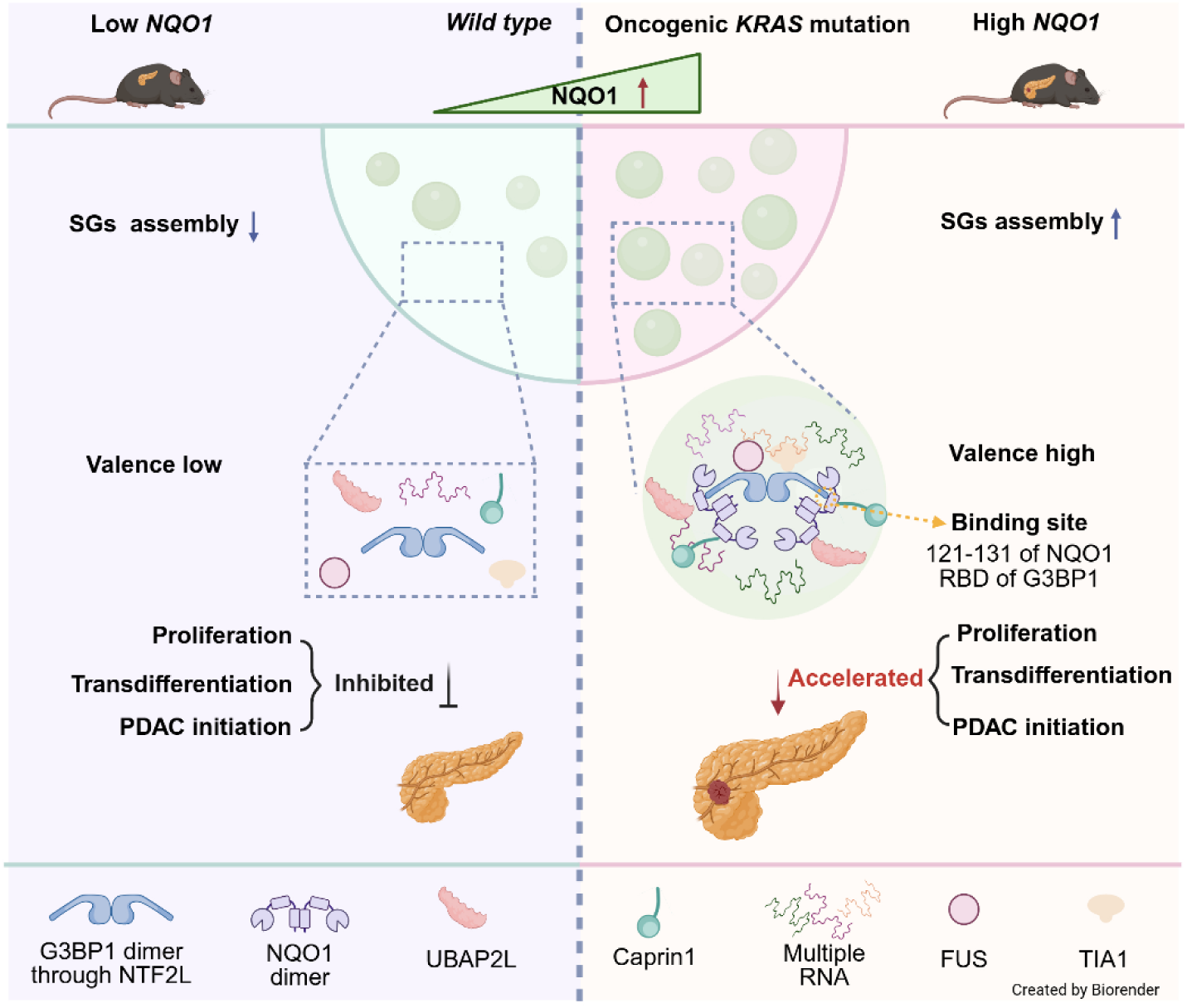
Schematic of KRAS-NQO1-G3BP1/SG axis regulates pancreatic carcinogenesis. The oncogenic KRAS mutation induces the expression of *NQO1* in the PDAC progression. NQO1 binding with G3BP1’s RBD domain through 121-131 residues to enhance the valence of G3BP1 complex. This interaction promotes the assembly of SG, thereby accelerating the transdifferentiation of acinar cells, cell proliferation and the initiation of PDAC. Figure created using BioRender.

## Discussion

*KRAS* mutation was the principal oncogenic driver of PDAC initiation and progression^36^, with ∼90% of PDAC cases exhibiting *KRAS* mutations^37^. Acinar cells, the predominant functional unit of the exocrine pancreas, exhibit marked plasticity through their transdifferentiation into duct-like cells via a process termed ADM^38^. While inflammation-induced ADM is typically reversible upon resolution of inflammatory stimuli, murine models harboring activating *Kras* mutations (e.g., *Kras^G12D^*) demonstrate irreversible ADM progression to PanIN and ultimately invasive PDAC^10, 39^. Despite its centrality to pancreatic oncogenesis, the molecular mechanisms underlying *KRAS*-driven ADM-PanIN-PDAC transition remain incompletely characterized.

Emerging evidence indicates that *KRAS* mutations orchestrate dysregulated assembly of SG. Spatial profiling in *Kras^G12D/+^*; *Trp53^R^*^172^*^H/+^*; *Pdx1^Cre^* (KPC) murine models and human PDAC tissues reveals predominant localization of G3BP1/SG in ADM and PanIN niches, suggesting an oncogenic role of SG during early PDAC pathogenesis^15^. Inhibition of SG results in higher cell death in *KRAS* mutant cells under oxidative stress compared to *KRAS* wild-type cells, suggesting that SG levels may be associated with the survival needs of *KRAS* mutants^15^. Additionally, in an obesity-induced PDAC model, SG formation is significantly upregulated. *G3BP1* gene knockdown or loss of its dimerization functional domain impedes SG formation, impairing PDAC proliferation, and preliminarily identifies SG as a key mediator in the pathophysiological function of PDAC^16^. Mechanistically, NUPR1—a key regulator of SG dynamics—promotes SG assembly, and pharmacological NUPR1 inhibition attenuates ADM/PanIN formation in KC mice^17^, further validating the functional centrality of SG in KRAS-driven pancreatic carcinogenesis^15^. While these studies delineate KRAS-SG interplay in PDAC progression, the precise molecular circuitry linking mutant KRAS to G3BP1/SG dysregulation remains incompletely resolved.

This study delineates NQO1 as a novel SG assembly modulator operating through ROS-independent mechanisms. While NQO1 has been characterized as a canonical antioxidant enzyme regulating oxidative stress responses in malignancies^8, 30, 31^, notably through ROS suppression to confer chemo-resistance^8^, this newly identified SG-regulatory role expands its functional repertoire. Previous RIP-seq data suggesting NQO1’s RNA-binding potential^25^, this study elucidates the structural and biophysical basis of this activity. NQO1 harbors a KH-type RNA recognition module featured with dual GXXG loops that colocalize with its IDR, and this KH domain is required for NQO1’s LLPS capacity. These findings establish NQO1 as a multifunctional hub coordinating RNA-protein condensate dynamics, thereby broadening its oncogenic mechanisms beyond redox regulation. We demonstrate that NQO1 directly bind with the RBD of G3BP1 via its 121-131 residues, synergistically enhancing G3BP1-driven LLPS and SG nucleation. Interestingly, the interaction between NQO1 and G3BP1 not only promotes G3BP1 LLPS but also required for NQO1 phase condensation during stress condition. The NTF2L and RBD domains of G3BP1 are mutually essential for LLPS and SG assembly. Previous studies showed that RNA binding proteins UBAP2L and Caprin1 increase SG complex valency (v=2→3) through NTF2L binding, thereby boosting G3BP1 LLPS. In contrast, USP10 lacks RBD and diminishes valency (v=2→ 1) via similar interactions, significantly impairing G3BP1 phase separation^23^. To date, few studies have reported RBD-dependent regulation of G3BP1 LLPS by SG regulators. Our findings demonstrate that NQO1’s 121-131 residues bind to G3BP1’s RBD, dramatically promoting phase separation at threshold concentrations. The pro-LLPS effect of NQO1 dependent on its scaffolding role: RBD binding exposes RNA-binding domains that recruit RNA and high valencies RBPs (e.g., UBAP2L/Caprin1/TIA1/FUS; Fig. 4 f,g), elevating SG complex valence to promote phase transition (Fig. 7). Our work advances the valence-based model of SG formation through characterization of RBD-dependent binding, offering fresh perspectives on SG assembly principles.

Emerging clinical and preclinical evidence reveals that oncogenic *KRAS* mutations drive *NQO1* upregulation^40^ (Fig. 5d), with elevated *NQO1* expression serving as a prognostic biomarker for adverse outcomes and accelerated PDAC progression^11^. In the meantime, Nrf2 was dysregulated in murine PDAC lesions and genetic ablation of *nrf2* attenuated the initiation of PDAC in animal models^4–6^. Given the well-established fact that Nrf2 transcriptional activates NQO1 expression^41^, it is reasonable to speculate that oncogenic *KRAS* mutations activates NRF2 to enhance *NQO1* expression in PDAC cells. Through a multimodal experimental framework— encompassing *in vitro* human PDAC cell lines, organoid cultures of primary acinar explants and genetically engineered KC mice—we delineated the mechanistic interplay between the KRAS-NRF2-NQO1-G3BP1/SG network and pancreatic carcinogenesis. Integrative analysis of human PDAC transcriptomic datasets further validated these findings, revealing weak association between *NQO1* and *G3BP1*. Importantly, we found both NQO1 and G3BP1 formed biological condensates and co-localized in the lesions of human PDAC tissue sections. Crucially, KRAS-driven *NQO1* overexpression amplifies SG assembly via the 121-131 residues binding G3BP1’s RBD, thereby elevating the valency of G3BP1 complexes to fuel pancreatic oncogenesis. Moreover, our findings of NQO1’s regulatory role on SG assembly further support the notion that NQO1 is the “quintessential” cytoprotective enzyme beyond ROS regulation^42, 43^. Oncogenic *KRAS* mutations hijack this cytoprotective function of NQO1 to facilitate pancreatic carcinogenesis. Taken together, our work identified a previously unrecognized molecular cascade linking KRAS-mediated ADM to phase transitions-driven SG assembly, proposing that NQO1-G3BP1 interaction may serve as a diagnostic biomarker and therapeutic target for early-stage PDAC intervention.

## Acknowledgements

We gratefully acknowledge Professor Ying Hu (Harbin Institute of Technology) and Professor Yong Liu (Xuzhou Medical University) for their constructive comments on this research. This work was supported by the National Natural Science Foundation of China (Grant No. 82203311,), the Basic Research Program of Jiangsu (Grant No. BK20220670), the Jiangsu Province Capability Improvement Project through Science, Technology and Education (NO. CXZX202234), the Open Project of Key Laboratory of Science and Engineering for the Multi-modal Prevention and Control of Major Chronic Diseases, Ministry of Industry and Information Technology (Grant No. MCD-2023-1-01), the Research Foundation of Xuzhou Medical University (D2023004, D2023012, D2022034), the Science and technology project of Xuzhou (Grant No.KC23003), the Fundamental Research Funds for the Central Universities (No. 2024QN11068), “Double Innovation” Doctor (No. 140924007), “Double Innovation” talents projects of Jiangsu Province, the Basic Science (Natural Science) Research Project of Higher Education Institutions in Jiangsu Province (24KJD320006). Jiangsu Postgraduate Research and Practice Innovation Program (KYCX24_3079 and KYCX25_3315). The funders had no roles in study design, data collection and analysis, decision to publish, or preparation of the manuscript.

## Author contributions

Conceptualization was the responsibility of WJ.G, X.L. Methodology was the responsibility of X.L, JK.D, YF.Z, Y.H, C.H, YL.C, N.W, RH.Y, M.W, ZJ.Z, LY.M, J.X, QD.L, YR.Z, HX.Z, WH.Z, JT.X. The investigation was conducted by WJ.G, X.L, JK.D, YF.Z, Y.H. The original draft was written by X.L and JK.D. The reviewing and editing were performed by WJ.G,J.B, Y.L, Y.H and KK.T. Fund acquisition was carried out by WJ.G, X.L, Y.H, J.B and Y.L. Supervision was the responsibility of WJ.G, JN.Z.

## Competing interests

The authors declare no competing interests.

## Extended Data

**Fig. s1 Related to Fig. 1.**
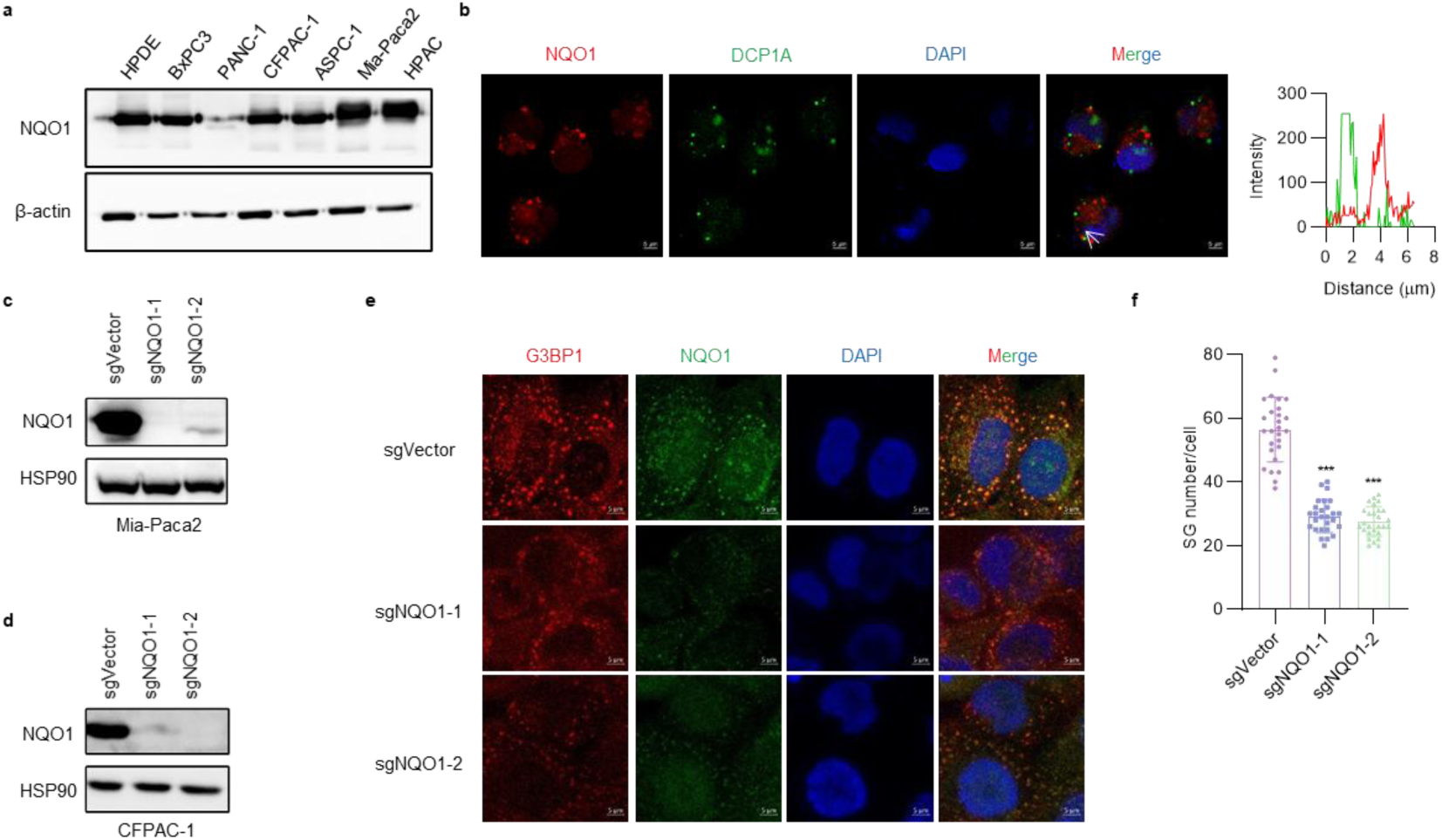
**a,** Expression of NQO1 in different PDAC cells. **b,** Confocal images of NQO1 clusters and P-bodies visualized by IF staining of NQO1 (red) along with DCP1A (green) in Mia-PaCa2 cells treated with NaAsO_2_ at 1μM for 1 h. Nucleus was stained with DAPI (blue). Scale bar, 5 μm. The white arrow indicates the colocalization and the fluorescence intensities is shown on the right. **c-d,** Western blot analysis was performed to validate the knockout efficiency of *NQO1* in *NQO1*-KO Mia-PaCa2 and CFPAC-1 cells. **e-f,** Following treatment with NaAsO_2_ (1 mM, 1 h), G3BP1 immunofluorescence staining was performed in Vector and *NQO1*-KO CFPAC-1 cells to assess SG formation. Quantitative analysis of SG numbers was conducted across three independent experiments (minimum 20 cells per experiment), and statistical differences were evaluated using unpaired Student’s t test. Mean ± s.d; ***, *p* < 0.001.

**Fig. s2 Related to Fig. 2.**
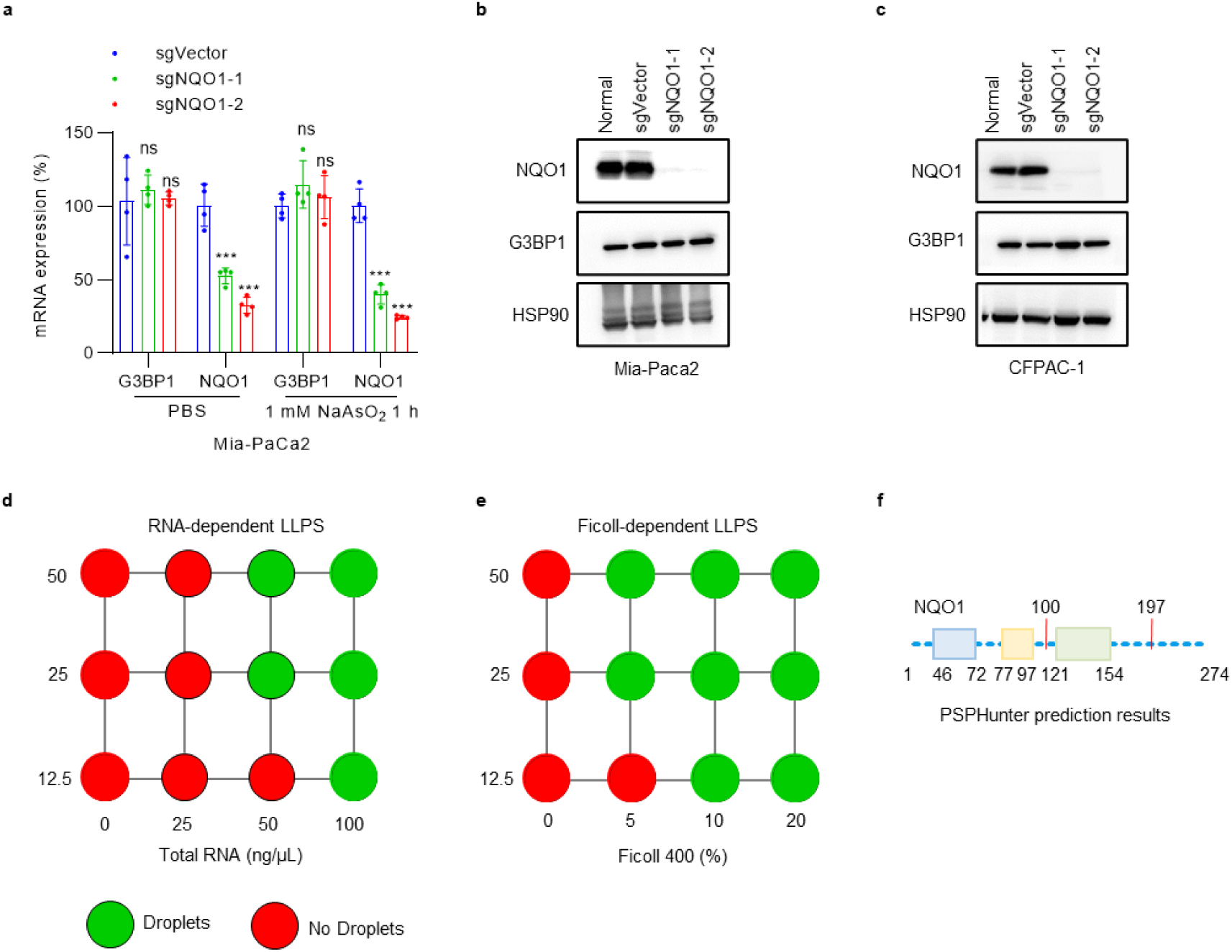
**a,**qRT-PCR analysis determined the regulatory role of *NQO1* knockout on *G3BP1* mRNA expression in Mia-PaCa2 cells under basal and stress-induced conditions. n=4; mean ± s.d; ns, not significant, *p* > 0.05; ***, *p* < 0.001. **b-c.** Western blotting revealed the impact of *NQO1* knockout on G3BP1 protein expression in Mia-PaCa2 and CFPAC-1 cells under physiological conditions. HSP90 was the protein loading control. **d-e.** Summary of phase separation behaviors of NQO1 in Fig2.d-e. **f,**PSPHunter predicts the key amino acid sites where NQO1 undergoes LLPS.

**Fig. s3 Related to Fig. 3.**
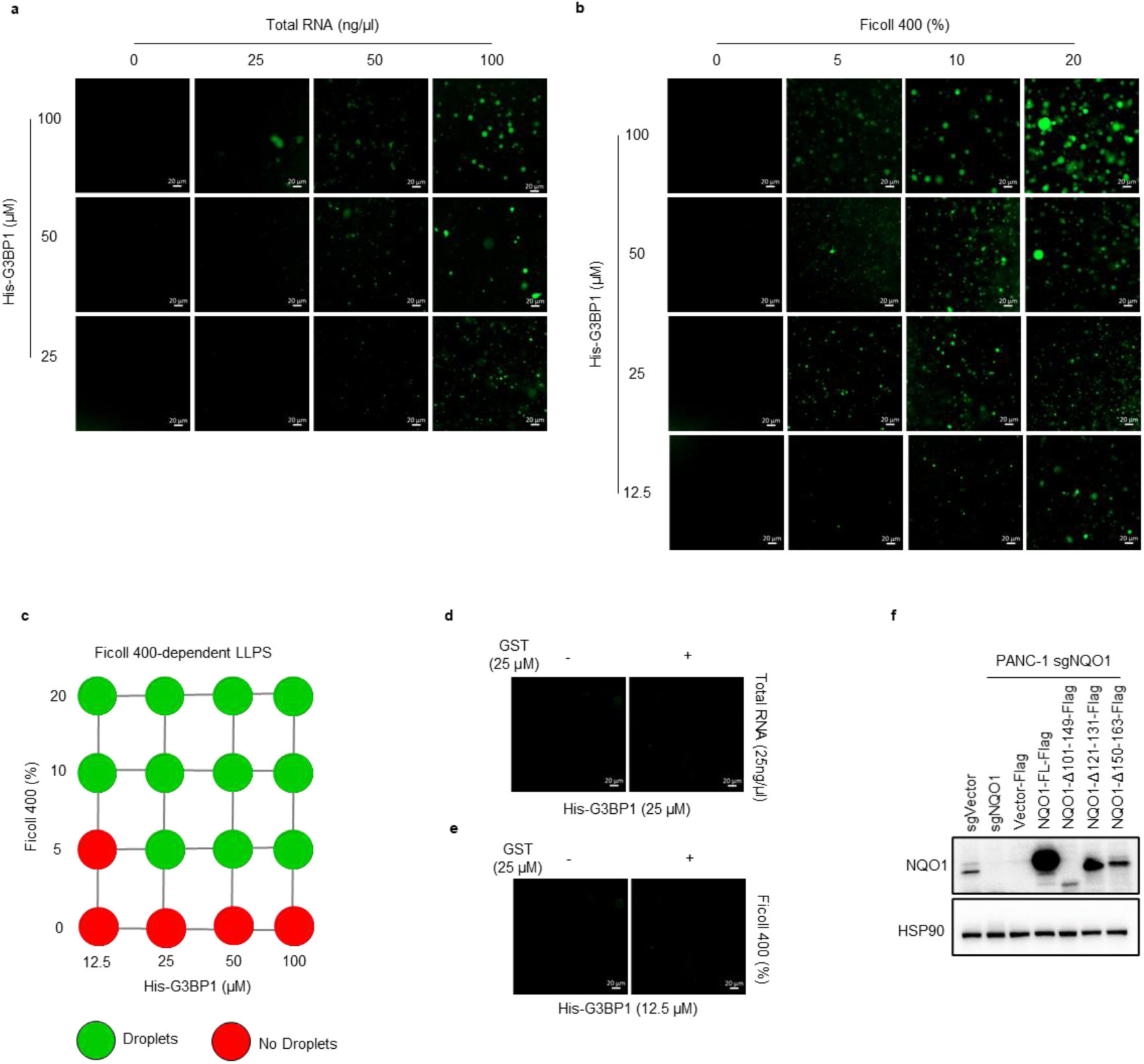
**a,** Representative images of phase separation of G3BP1 FL protein under varying concentrations of Total RNA. Scale bar, 20 μm. **b,** Characteristic phase separation behaviors of G3BP1 FL protein exposed to varying Ficoll 400 concentrations. Scale bar, 20 μm. **c,** Comprehensive analysis of G3BP1 phase separation properties derived from Fig. s3b. **d-e,** NHS-488 ester-stained G3BP1 full-length protein demonstrating phase separation characteristics upon GST protein addition at critical phase transition concentrations. Total RNA-containing versus RNA-free environments (d), Ficoll 400-supplemented and Ficoll-deficient conditions (e). Scale bar, 20 μm. **f,** *NQO1* KO PANC-1 cell transfected with pCDNA3.1-Flag-NQO1 FL and NQO1 truncations, western blot detect the expression. HSP90 was the protein loading control.

**Fig. s4 Related to Fig. 4.**
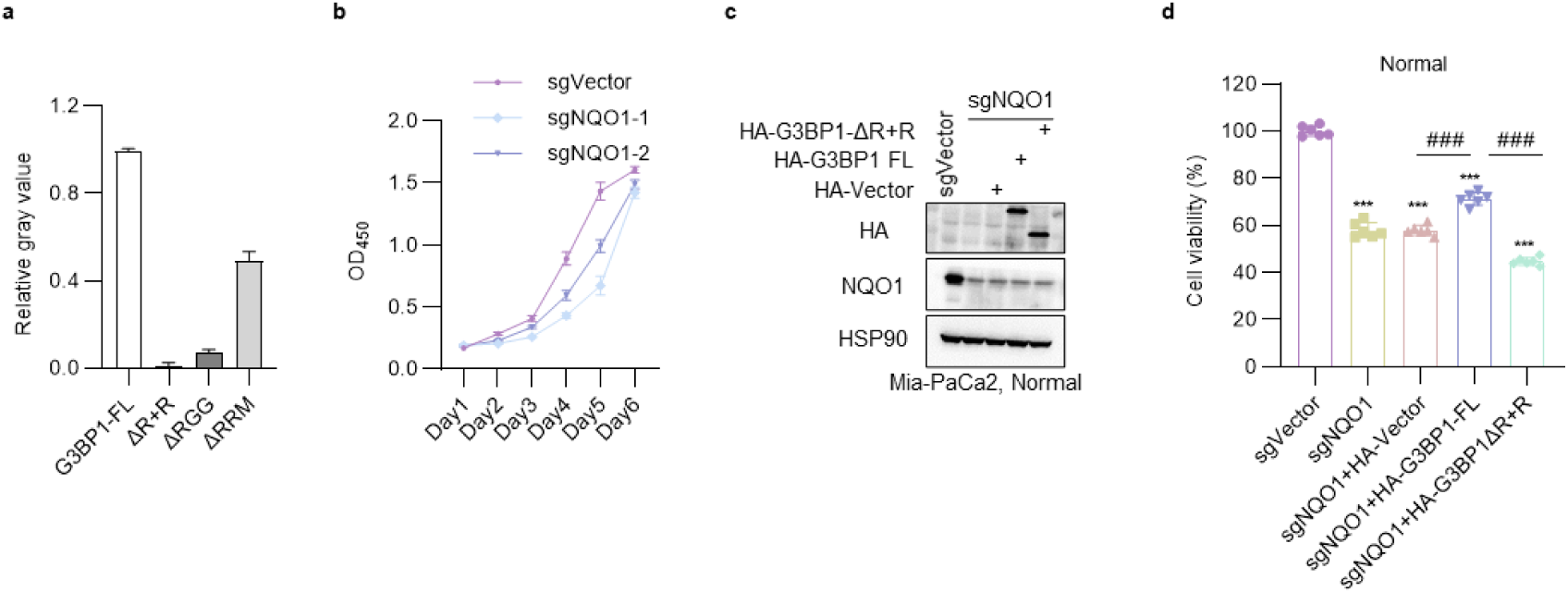
**a,** Statistics of protein gray values as a result of two experiments in Fig. 4d. Data are represented as mean ± s.d. **b,** Proliferative capacity of *NQO1*-KO Mia-PaCa2 cells assessed by CCK-8 assay. **c,** The expression levels of PKH3-HA-G3BP1 FL and PKH3-HA-G3BP1-ΔR+R in *NQO1* KO Mia-PaCa2 cells under normal conditions, the expression were determined by Western blot analysis. HSP90 was the protein loading control. **d,** Cell proliferation assays of *NQO1* KO Mia-Paca2 cells transfected with PKH3-HA-Vector, PKH3-HA-G3BP1 FL, or PKH3-HA-G3BP1-ΔR+R under normal conditions (n=6 independent experiments). Data were analyzed using unpaired Student’s t test. ***, *p* < 0.001; compared to sgVector; ^###^, *p* < 0.001; relative to sgNQO1+HA-G3BP1 FL group.

**Fig. s5 Related to Fig. 5.**
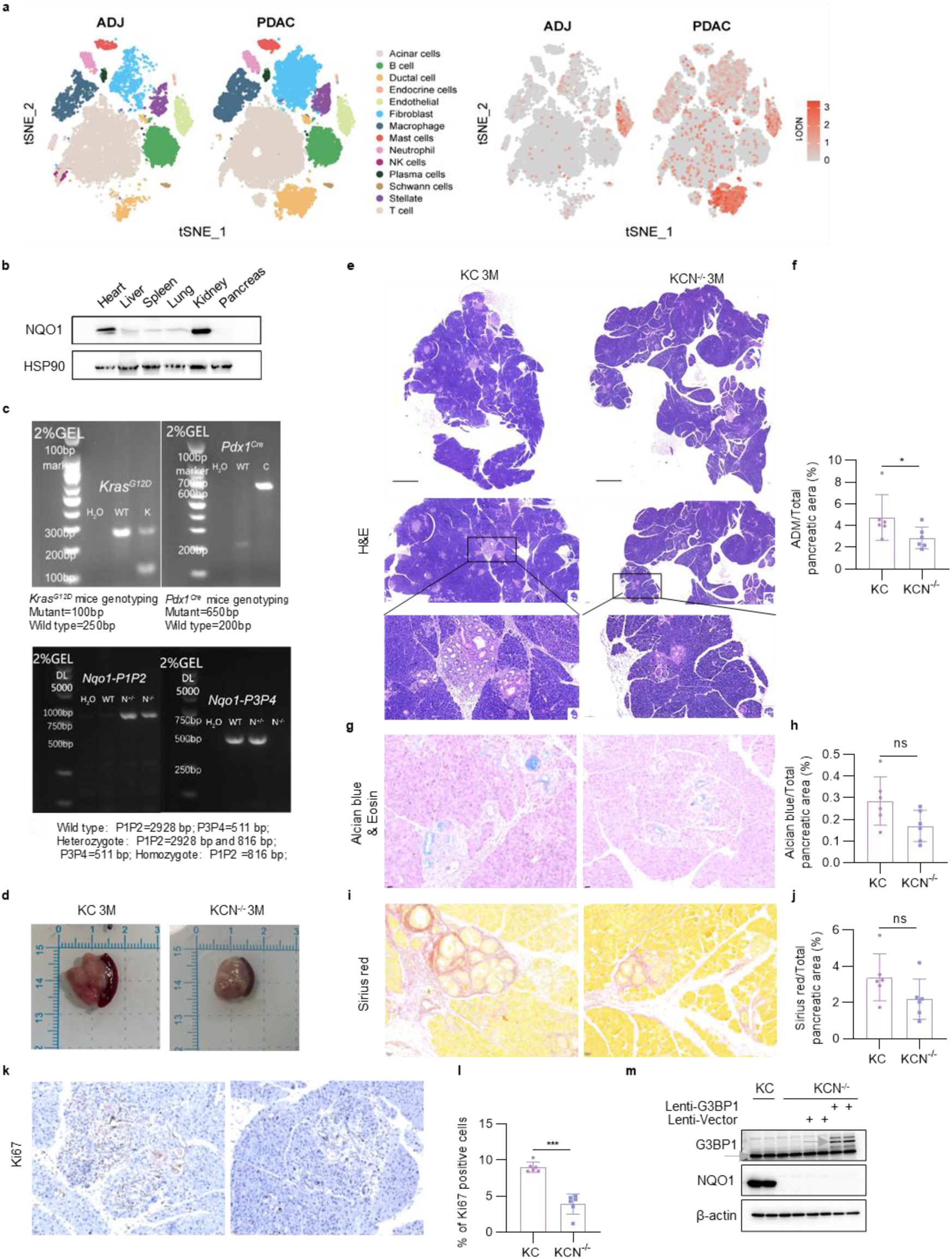
**a,** Analysis of single-cell sorting results of *NQO1* in human pancreatic tumor and normal tissues. **b,** NQO1 protein expression in wild-type mouse organs was validated by Western blotting.HSP90 was the protein loading control. **c,** PCR-based genotyping verification of KC and KCN^-/-^ mice. **d,** Morphological presentation of pancreatic tissues from 3-month-old KC and KCN^-/-^ mice. **e,** Representative H&E-stained pancreatic tissue sections from 3-month-old KC and KCN^-/-^ mice (2×, 10×, and 20× magnifications; scale bars: 1000 μm, 200 μm, and 100 μm respectively). **f,** Statistical analysis of ADM area proportion relative to total pancreatic area using H&E staining results. **g-h,** Alcian Blue/Eosin staining and statistical analysis of Alcian Blue-positive regions in total pancreatic area. **i-j,** Sirius Red staining with statistical evaluation of collagen-positive area proportion. **k-l,** Ki67 IHC staining with statistical analysis of proliferating cell ratio. Results in **e-l** represent pancreatic sections from 3-month-old KC and KCN^-/-^ mice (n=6). Intergroup differences were assessed by unpaired Student’s t test (n=6; data shown as mean ± s.d). **, *p* < 0.01; ***, *p* < 0.001; ns, not significant, *p* > 0.05. **m,** G3BP1 expression verification in primary acinar cells post-lentiviral plasmid delivery (Fig. 5s experiment).

## Materials

### Methods

#### Cell culture and stable cell line construction

Human pancreatic cancer cells (BxPC3, ASPC-1, HPAC, PANC-1, Mia-PaCa2 and CFPAC-1) were obtained from the Cell Resource Center, Institute of Basic Medical Sciences. HEK-293T cells was gifted by the laboratory of Prof. Liu Yong, Xuzhou Medical University. All cell lines were cultured in DMEM or RPMI 1640 medium (C11995500BT, Gibco), supplemented with 10% fetal bovine serum (ST30-3302, PAN-Biotech GmbH) and 1% penicillin/streptomycin solution (VC2003, VICMED). Cells were maintained at 37 °C in a humidified incubator with 5% CO_2_. Mycoplasma contamination was assessed using the Mycoplasma PCR Detection Kit (C0301S, Beyotime), and cell line authenticity was confirmed by STR profiling (Wuhan Pricella Biotechnology).

Lentiviral particles were generated in HEK-293T cells by co-transfecting the lentiviral plasmid along with packaging plasmids pMD2.G and pSPAX2 using Lipo8000 Transfection Reagent (C0533, Beyotime). 48 hours post-transfection, the culture supernatant containing the lentivirus was harvested, filtered through a 0.22 μm filter, and used to infect target cell lines. Two days after infection, the medium was replaced with puromycin-containing medium for selection. Single-cell clones were isolated by limiting dilution in 96-well plates and expanded for further analysis.

#### Plasmids and transfection

Plasmid information is detailed in Table 3. All plasmids listed in the table were constructed using either Gibson assembly or homologous recombination, with the exception of those explicitly marked with their source. Gibson assembly was conducted following PCR amplification (Q5 Hot Start High-Fidelity 2× Master Mix, M0494S, NEB). Truncated and mutant forms of NQO1 and G3BP1 were generated via KLD substitution (M0554S, NEB). The TIA1 fragment was cloned from pCMV-TIA1(human)-Neo into the PKH3-HA-Vector, employing the same methodology for UBAP2L, Caprin1, and FUS (Purchased at MIAOLING PLASMID). All sequences were subsequently verified through Sanger sequencing. Primer sequences are available as Supplementary Data.

**Table1.**
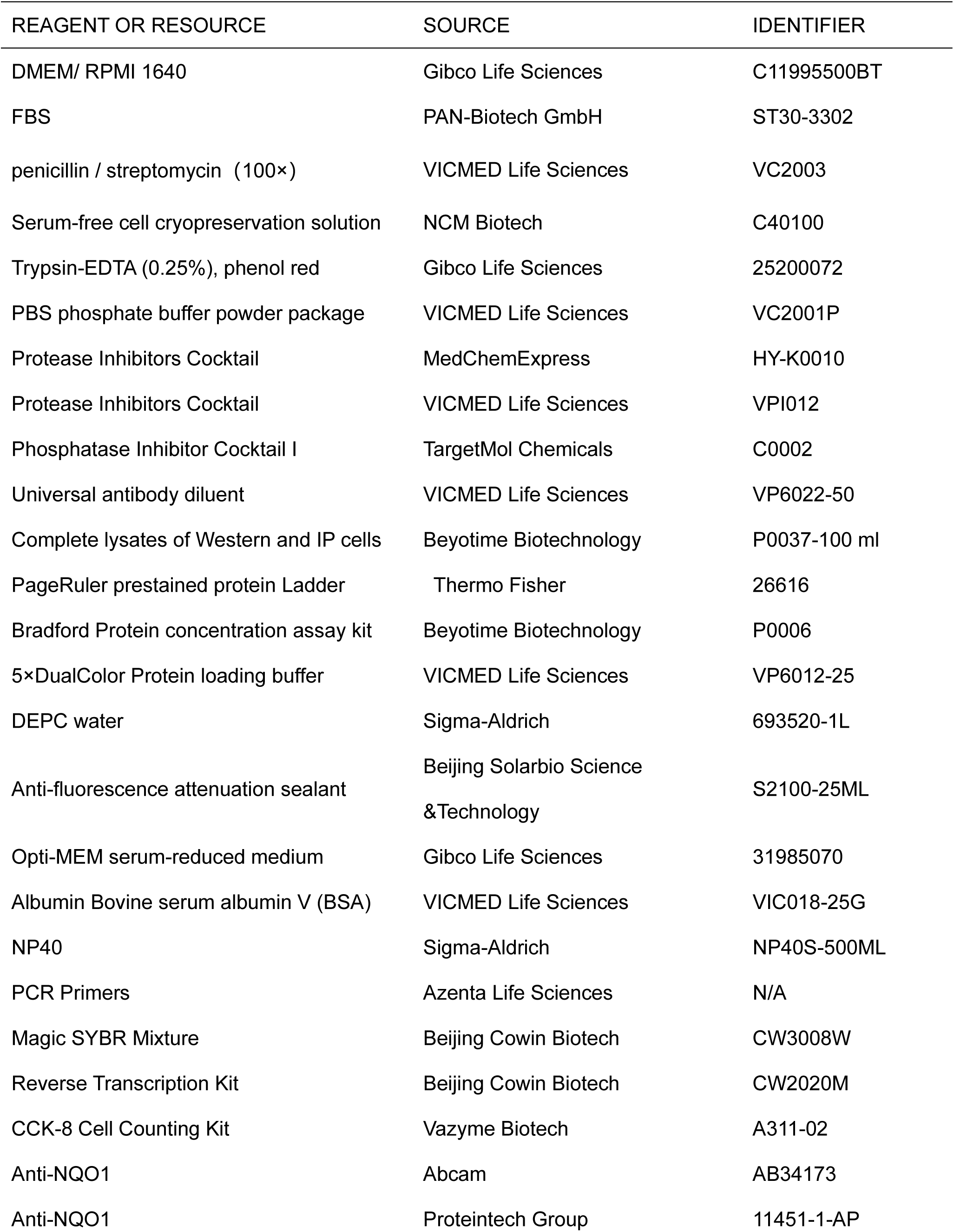
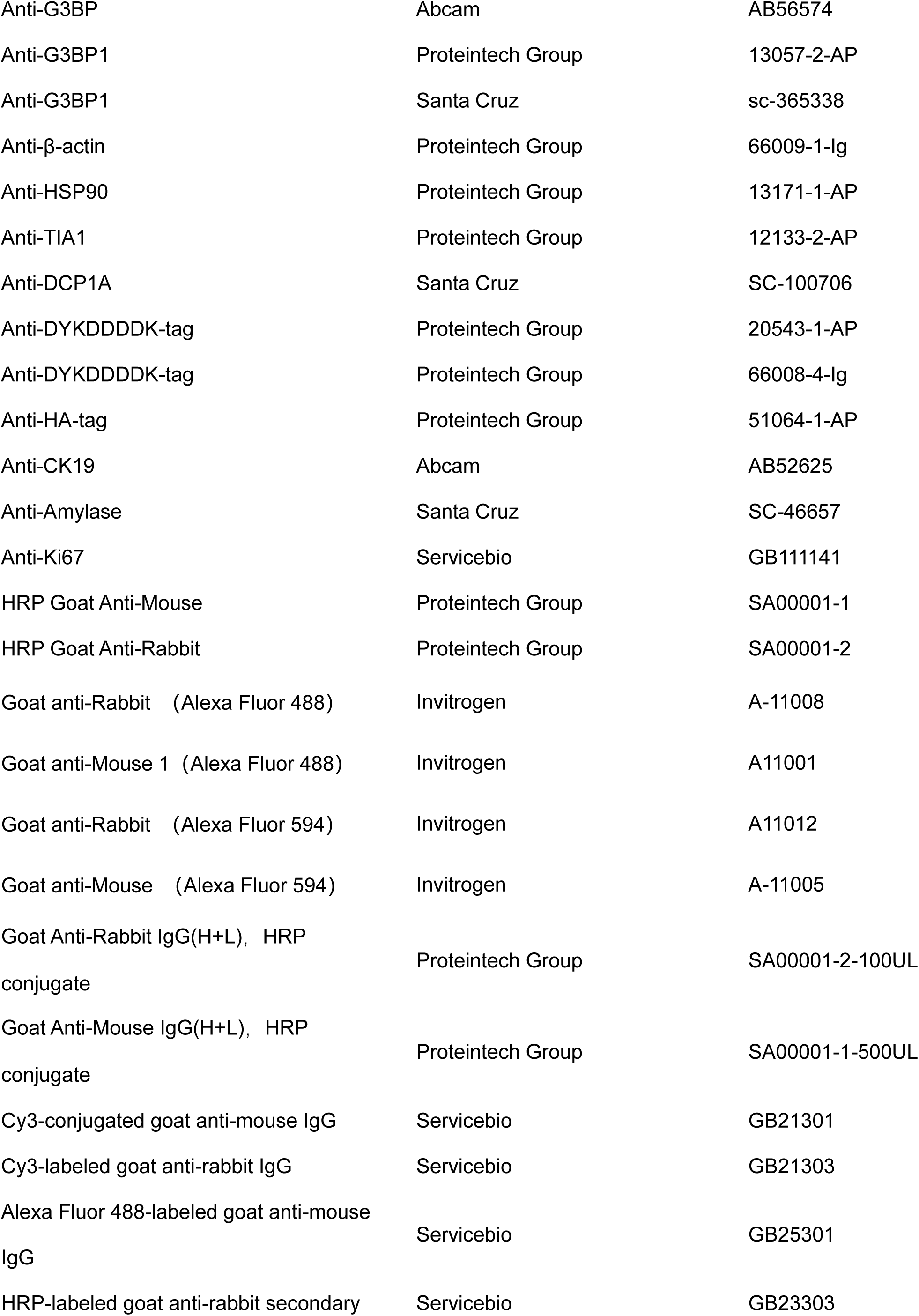
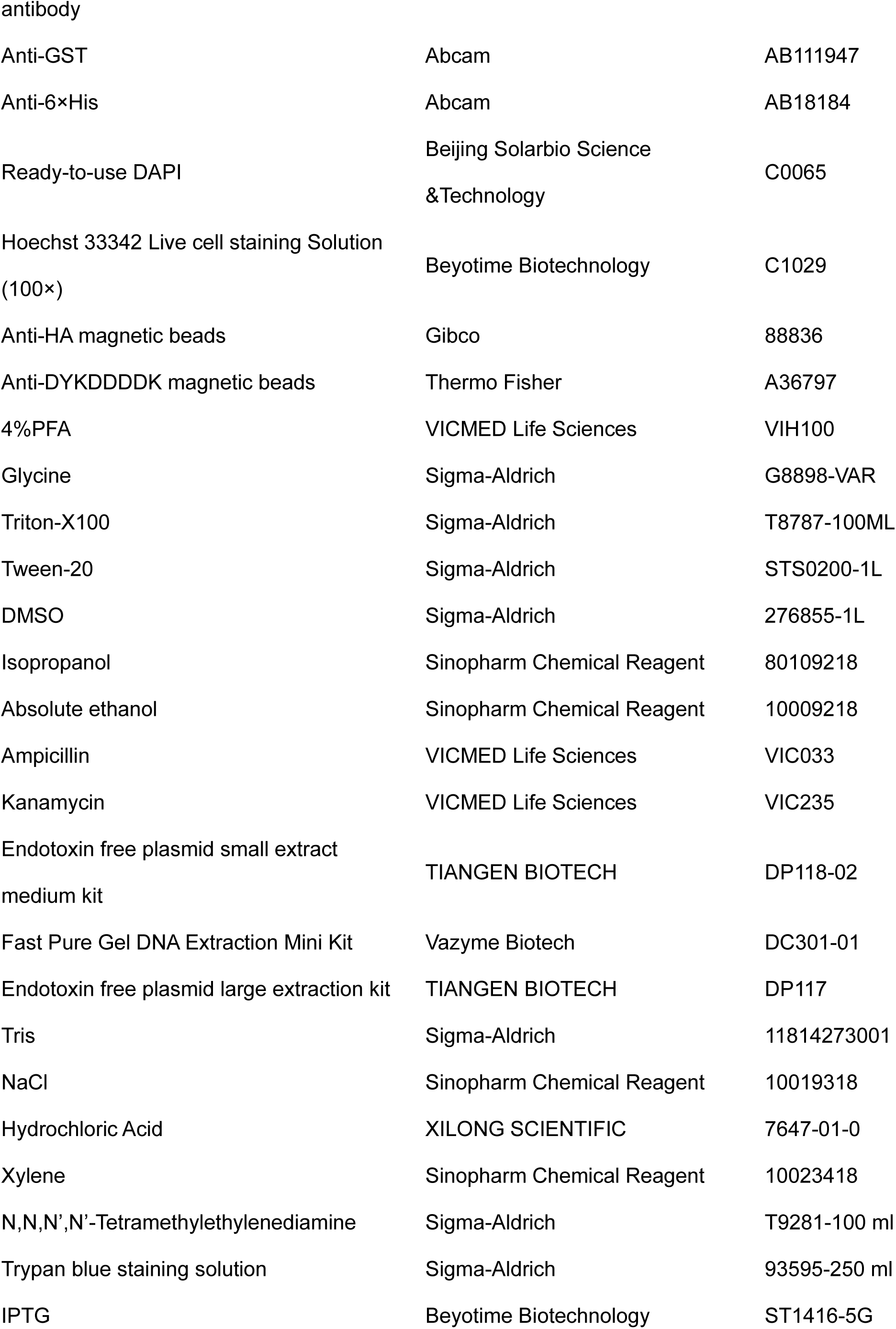
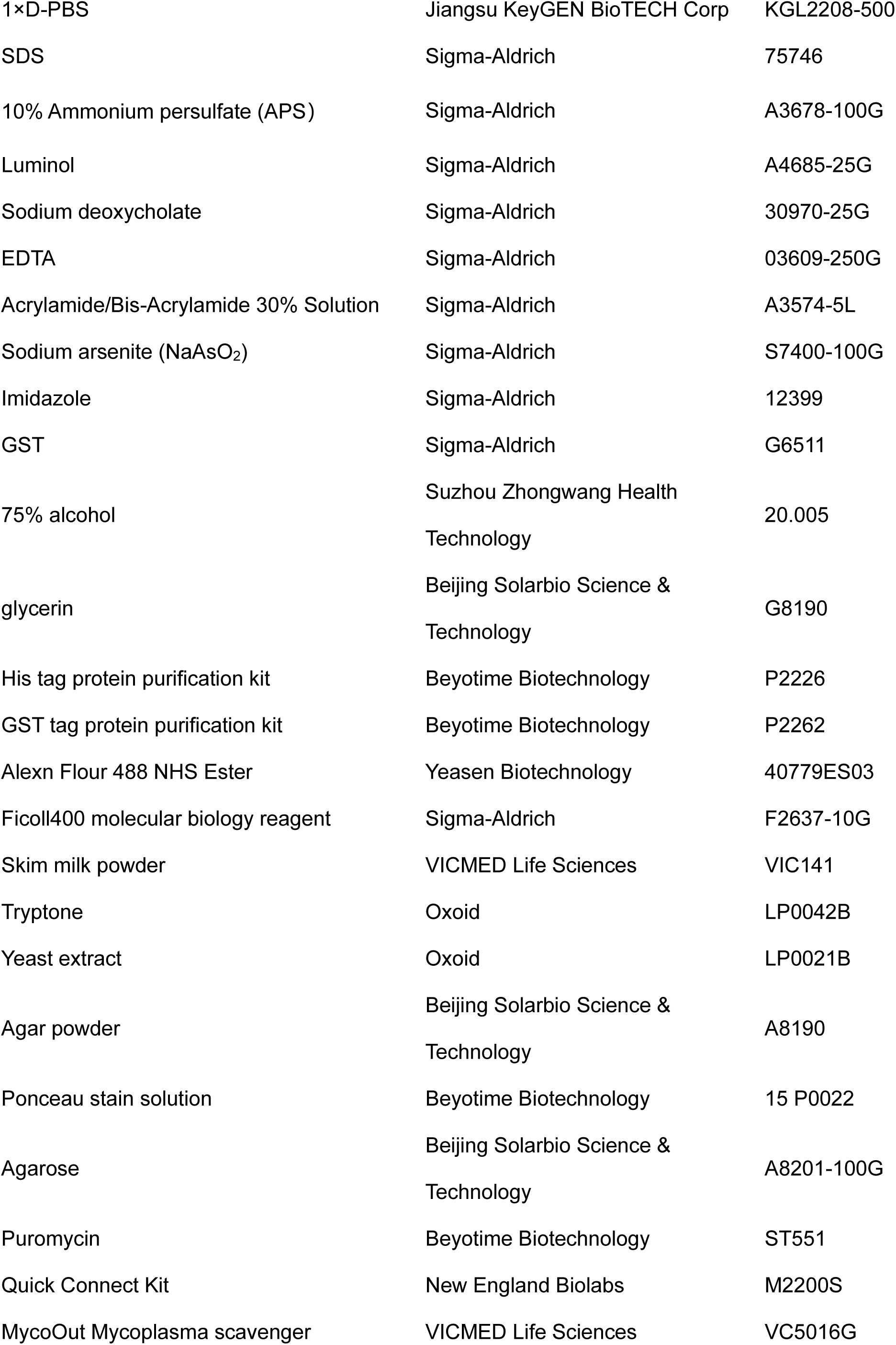
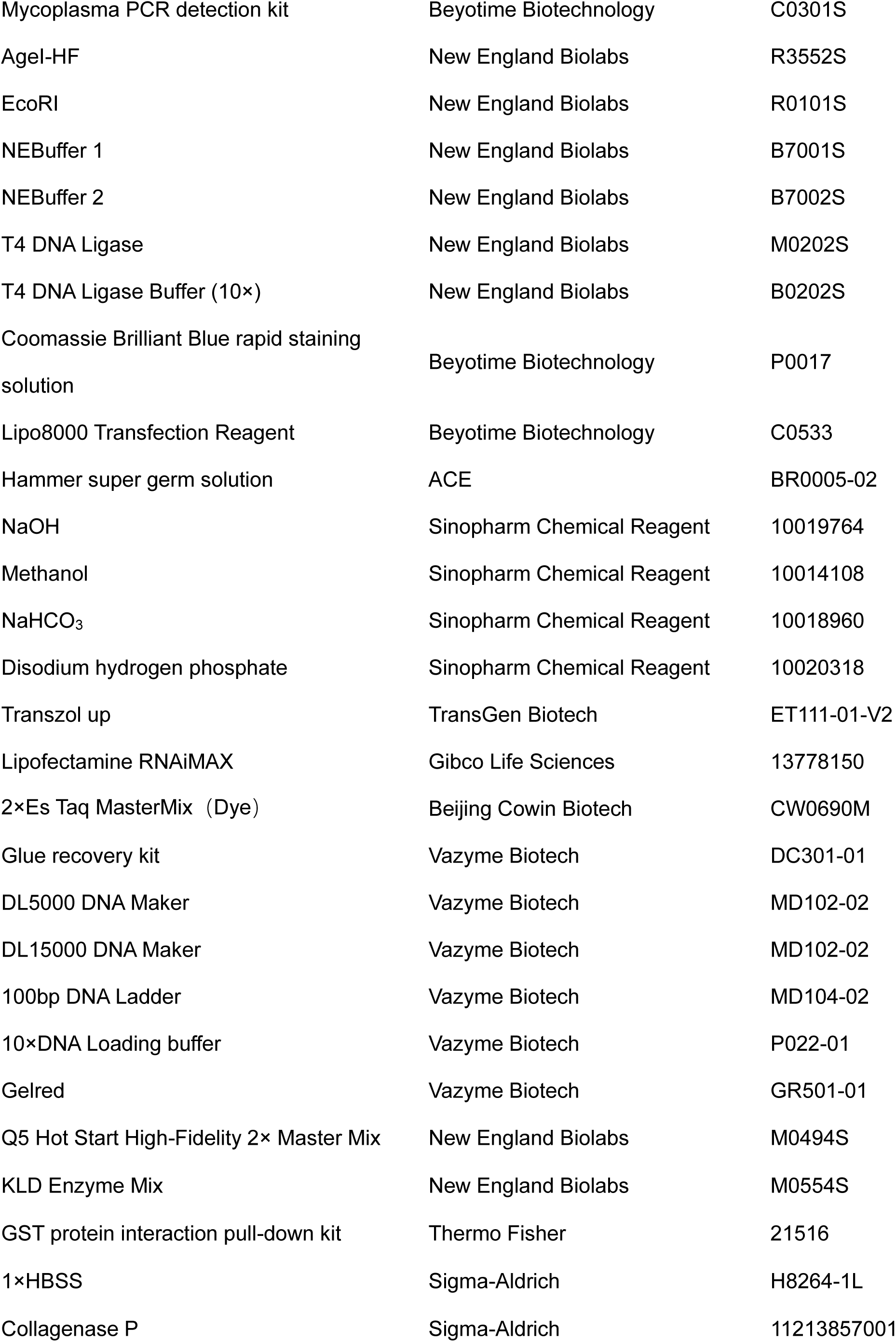
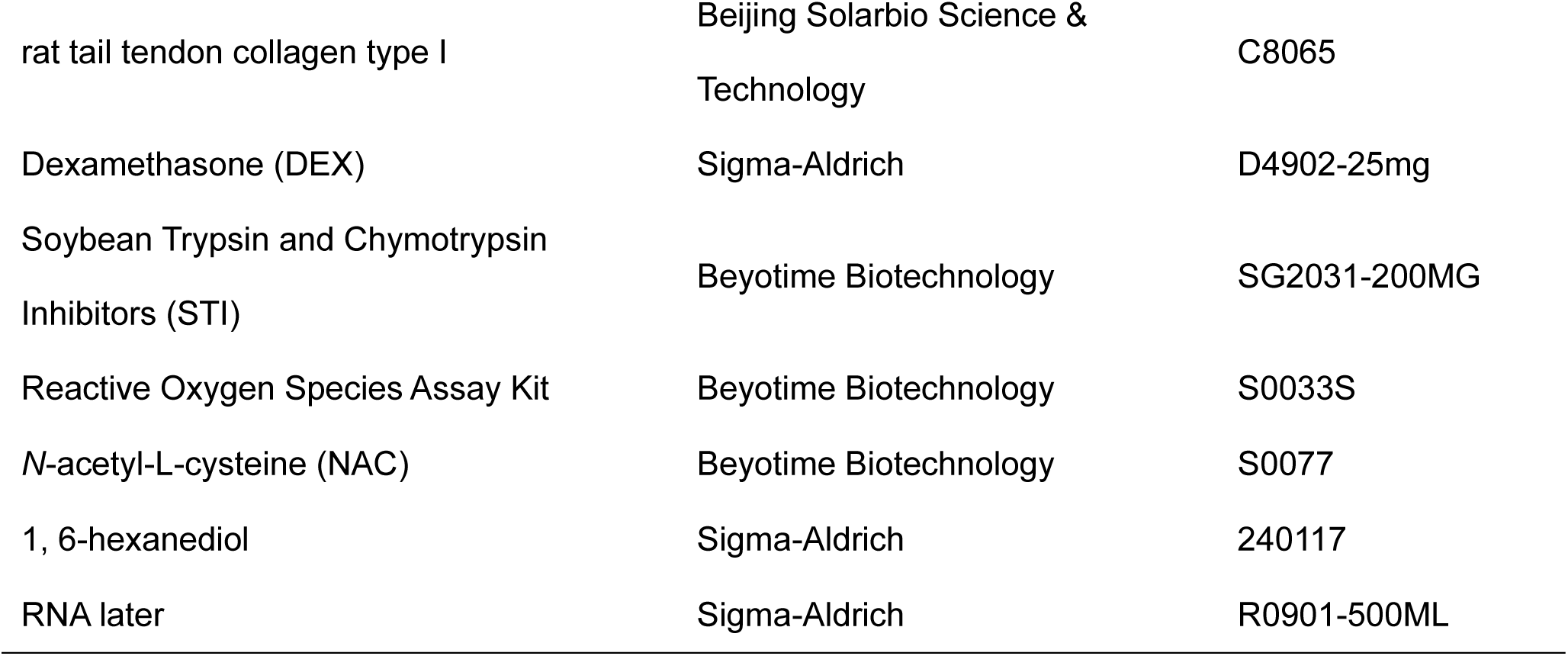
Primary Reagent.

**Table2.** Recombinant DNA.

| PLASMID | SOURCE |
| --- | --- |
| Lenti-Crispr V2 | Unibio |
| psPAX2 | Unibio |
| pMD2.G | Unibio |
| Lenti-CRISPR V2 sgNQO1-1 | This study |
| Lenti-CRISPR V2 sgNQO1-2 | This study |
| pcDNA3.1-3xFlag | Zhang Yanping Laboratory, North Carolina |
| pcDNA3.1-3xFlag-NQO1 | Addgene(#61729) |
| pcDNA3.1-3xFlag-NQO1-FL-myc | This study |
| pcDNA3.1-3xFlag-NQO1-△1-100-myc | BioMed |
| pcDNA3.1-3xFlag-NQO1-△101-197-myc | BioMed |
| pcDNA3.1-3xFlag-NQO1-△198-274-myc | BioMed |
| pcDNA3.1-Flag-NQO1-△101-149 | This study |
| pcDNA3.1-Flag-NQO1-△150-163 | This study |
| pcDNA3.1-Flag-NQO1-△164-197 | This study |
| pcDNA3.1-Flag-NQO1-△101-110 | This study |
| pcDNA3.1-Flag-NQO1-△111-120 | This study |
| pcDNA3.1-Flag-NQO1-Δ121-131 | This study |
| pcDNA3.1-Flag-NQO1-Δ132-139 | This study |
| pcDNA3.1-Flag-NQO1-Δ140-149 | This study |
| PKH3-HA-Vector | This study |
| PKH3-HA-G3BP1-FL | This study |
| PKH3-HA-G3BP1-ΔRGG | This study |
| PKH3-HA- G3BP1-ΔR+R | This study |
| PKH3-HA- G3BP1-ΔR+R+P | This study |
| PKH3-HA- G3BP1-ΔR+R+P+A | This study |
| HA- G3BP1-ΔRRM | This study |
| pGEX-4T-1-NQO1-GST | This study |
| pET-28a-G3BP1-His | This study |
| pET-28a-NQO1-His | This study |
| plv3-cmv-EGFP-NQO1-puro | This study |
| Opto-pHR-mCh-Cry2WT | Addgene (#101221) |
| Opto-NQO1-FL | This study |
| Opto-NQO1-Δ1-100 | This study |
| Opto-NQO1-Δ101-197 | This study |
| OptoNQO1-Δ198-274 | This study |
| PKH3-3xHA-FUS | This study |
| PKH3-3xHA-TIA1 | This study |
| PKH3-3xHA-UBAP2L | This study |
| PKH3-3xHA-CAPRIN1 | This study |
| pLV3-CMV-UBAP2L-3xFLAG-CopGFP-Puro | MiaoLing Biology |
| pCMV-CAPRIN1-3xFLAG-Neo | MiaoLing Biology |

**Table 3.**
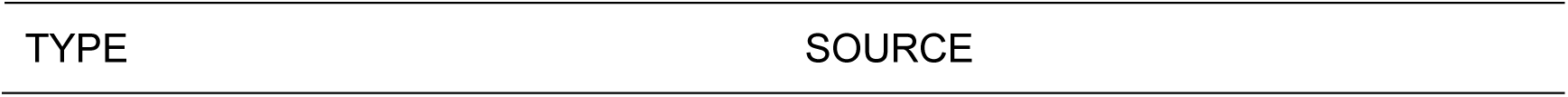
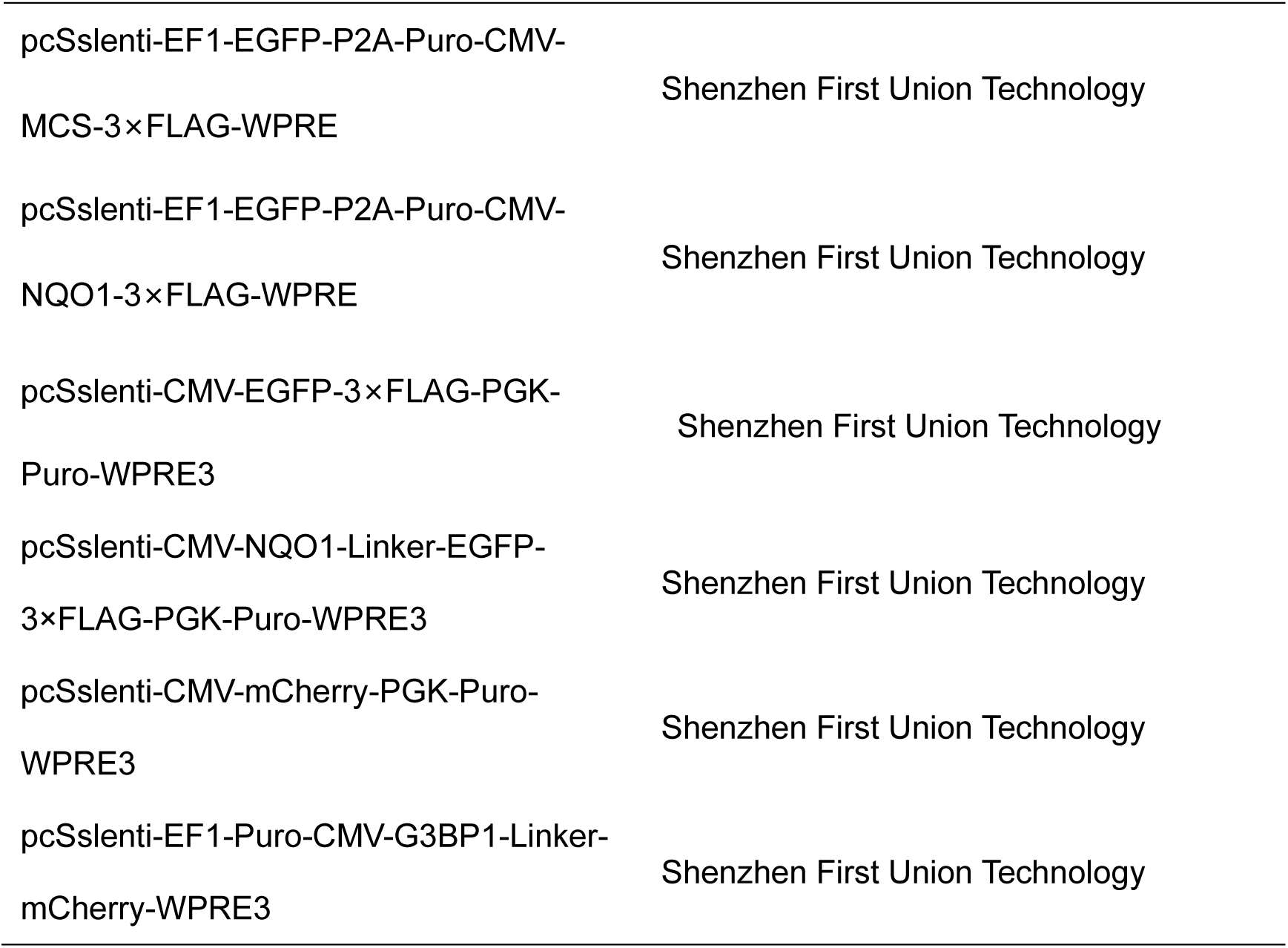
Virus strains.

| TYPE | SOURCE |
| --- | --- |
| pcSslenti-EF1-EGFP-P2A-Puro-CMV-MCS-3×FLAG-WPRE | Shenzhen First Union Technology |
| pcSslenti-EF1-EGFP-P2A-Puro-CMV-NQO1-3×FLAG-WPRE | Shenzhen First Union Technology |
| pcSslenti-CMV-EGFP-3×FLAG-PGK-Puro-WPRE3 | Shenzhen First Union Technology |
| pcSslenti-CMV-NQO1-Linker-EGFP-3×FLAG-PGK-Puro-WPRE3 | Shenzhen First Union Technology |
| pcSslenti-CMV-mCherry-PGK-Puro-WPRE3 | Shenzhen First Union Technology |
| pcSslenti-EF1-Puro-CMV-G3BP1-Linker-mCherry-WPRE3 | Shenzhen First Union Technology |

**Table 4.**
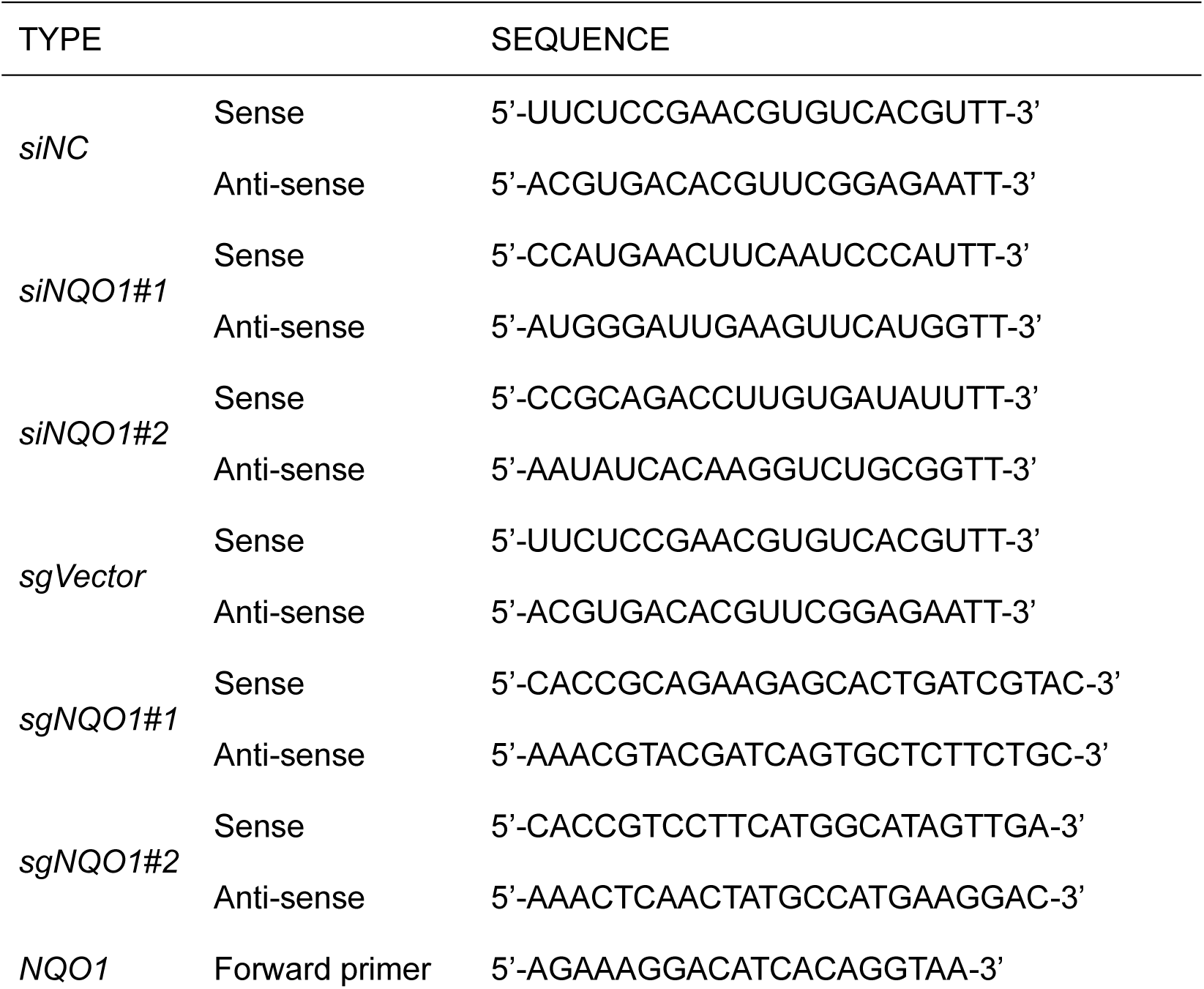
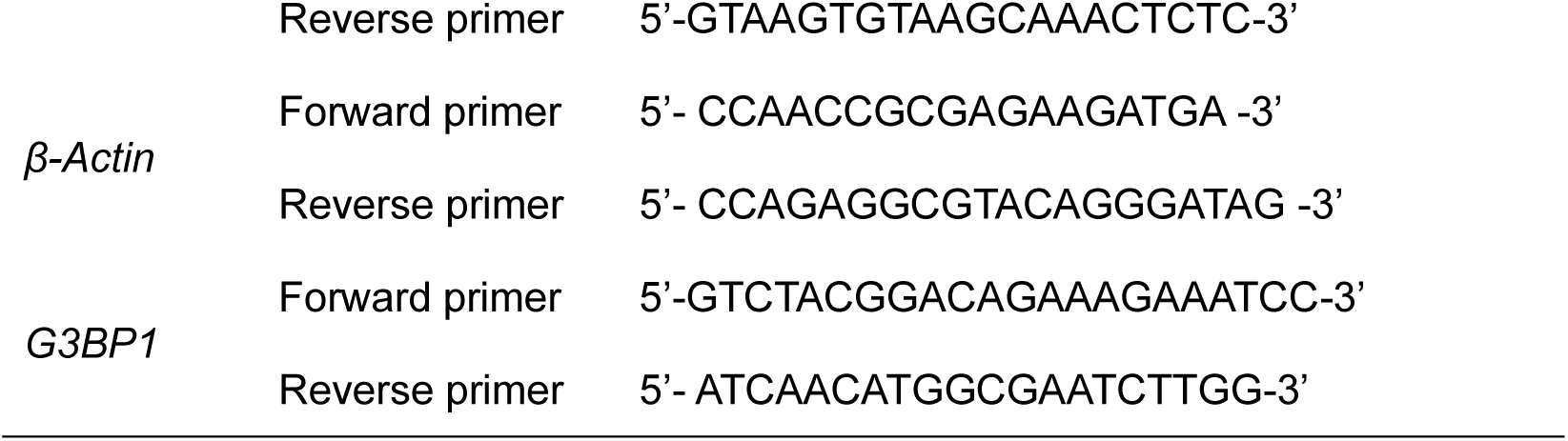
Sequence.

In 6-well plates, cells were cultured at a density of 30-40%. For transfection, each well received a mixture of 2 μg plasmid DNA, 2 μl Lipo8000 Transfection Reagent (C0533, Beyotime), and 200 μl Opti-MEM (31985070, Gibco). Prior to addition, 2 μg plasmid DNA was combined with 2 μl Lipo8000 in 100 μl Opti-MEM, incubated at room temperature for 20 minutes, and then added dropwise to the wells in the 6-well plates. Following 6-8 hours of incubation, the medium was replaced with fresh culture medium. Subsequent experiments were conducted 48 hours after transfection.

When performing siRNA transfection, each well was treated with a transfection mixture containing 50 nM siRNA, 2.5 μl Lipofectamine RNAiMAX (13778150, Gibco), and 200 μl Opti-MEM (31985070, Gibco). Prior to administration, 5 μl siRNA (20 μM) was complexed with 2.5 μl Lipofectamine RNAiMAX in 100 μl Opti-MEM, followed by a 20 minutes incubation at room temperature. The prepared complex was then added dropwise to the wells of 6-well plates. After 6-8 hours of incubation, the transfection medium was replaced with fresh culture medium.

#### Opto-Droplet detection experiment

The full-length NQO1 and truncated NQO1 fragments were fused with the pHR-mCh-Cry2WT (Opto, Addgene #101221) vector via homologous recombination. Subsequently, HEK-293T cells were seeded in confocal dishes, and the plasmids, including pHR-mCh-Cry2WT, Opto-NQO1-FL, Opto-NQO1-Δ1-100, Opto-NQO1-Δ101-197, OptoNQO1-Δ198-274 were transiently transfected using Lipo8000. After 48 hours of incubation, live-cell imaging was performed on a Live Cell Workstation. Cells were irradiated with a 488 nm laser for 1 second to activate Cry2. Fluorescence images were acquired at 10 seconds intervals over a period of 3 minutes in the mCherry channel. This cycle was repeated five times to ensure data consistency and reliability.

#### Single-cell data analysis

Single-cell expression profiles of PDAC and adjacent noncancerous tissues (GSE212966) were downloaded from GEO (https://www.ncbi.nlm.nih.gov/geo/). The quality control and downstream analysis of single cell RNA-seq data were performed by the R toolkit Seurat (v5.2.1)^44^. Low-quality cells with mitochondrial gene ratios exceeding 25%, total gene counts less than 500, and total gene expression levels below 1000 were removed. The R package DoubletFinder (v2.0.3)^45^ was used to eliminate doublet cells. Subgroup analysis of cells was conducted using marker genes as described in previous studies^46^, and the expression of NQO1 in PDAC and adjacent noncancerous (ADJ) cells was demonstrated.

#### RNA-seq data analysis

Following 1 h exposure to 1 mM sodium arsenite (NaAsO2) under standard culture conditions, both vector-control and NQO1-KO Mia-PaCa2 cells underwent two ice-cold PBS washes prior to Trizol-based lysis and subsequent cryopreservation at -80°C. RNA integrity evaluation and quantification services were provided by Novogene Corporation using Agilent Bioanalyzer 2100 system. FastQC was used to check reads quality and fastp^47^ removed low-quality reads. Reads were then mapped to human genome (hg38) with STAR^48^. RSEM^49^ was served to generate expression count files. Differentially expressed genes were identified by DESeq2^50^ R package, and selected genes with adjust p-value ≤0.05 and |log2(fold change)|≥1.

#### Mass spectrometry

Plasmid-mediated Flag-tagged NQO1 overexpression in Mia-PaCa2 cells was achieved through 48-hour transfection under optimal conditions. Three sequential PBS washes preceded cell disruption in NETN lysis buffer containing complete EDTA-free protease inhibitor cocktail (MCE, HY-K0010). Flag-tagged protein complexes were immune-precipitated with ANTI-FLAG beads (Thermo Fisher, A36797) following manufacturer’s protocol. Protein lysates were incubated with affinity matrix for 2 h at 4°C under constant agitation, with subsequent five-stage washing in high-salt NETN buffer. Protein denaturation and release from beads was achieved through 5-minute incubation at 95°C in reducing SDS-PAGE loading buffer (62.5 mM Tris-HCl, pH 6.8, 2% SDS, 10% glycerol). In-solution trypsinization was performed on purified proteins. LC-MS/MS analysis and raw data processing were conducted by Bioprofile Biotechnology using Q Exactive™ HF-X system. Gene Ontology (GO) enrichment analysis was performed using DAVID.

### RNA extraction and Real time quantitative reverse transcription-PCR (qRT-PCR)

Total RNA was isolated from cell samples using Transzol Up reagent (ET111-01-V2, TransGen Biotech) in strict adherence to the manufacturer’s protocol. The isolated RNA was then reverse transcribed into complementary DNA (cDNA) using a commercially available reverse transcription kit (CW2020M, Cowin), followed by real-time quantitative PCR (RT-qPCR) analysis with Magic SYBR Mixture (CW3008W, Cowin). Quantitative real-time PCR (qRT-PCR) analysis was conducted using the 2^-ΔΔCT^ method, with β-Actin serving as the internal reference gene for normalization. Relative gene expression levels were determined by normalizing to control samples. Data are presented as mean ± standard deviation (s.d) from three independent biological replicates. The specific primer sequences used for qRT-PCR analysis are detailed in Table 5.

### Immunofluorescence

For immunofluorescence analysis, cells were cultured on 24-well plates equipped with climbing sheets. Following the specified treatments, cells were fixed with 4% paraformaldehyde (PFA; VIH100, VICMED Life Sciences) at room temperature for 15 minutes and subsequently permeabilized with 0.4% Triton X-100 (T8787, Sigma) at 4°C for 5 minutes. Samples were blocked with 3% bovine serum albumin (BSA; VIC018, VICMED Life Sciences) for 1 hour at room temperature, followed by overnight incubation with specific primary antibodies targeting the proteins of interest at 4°C. Slides were washed three times with phosphate-buffered saline (PBS; 10 minutes each) and then incubated with secondary antibodies at room temperature for 2 hours. Subsequently, slides were stained with either 4’,6-diamidino-2-phenylindole (DAPI; C0065, Solarbio) or Hoechst 33342 Live Cell Staining Solution (C1029, Beyotime Biotechnology) at room temperature for 15-30 minutes, followed by three additional PBS washes (10 minutes each). Finally, the climbing slides were mounted using an anti-fluorescence quenching mounting solution (S2100, Solarbio).

The following antibodies were utilized: Goat anti-Mouse IgG (H+L) Alexa Fluor 594 (A-11005, Invitrogen) at 1:200 dilution, Goat anti-Rabbit IgG (H+L) Alexa Fluor 488 (A-11008, Invitrogen) at 1:200 dilution, Goat anti-Rabbit IgG (H+L) Alexa Fluor 594 (A11012, Invitrogen) at 1:200-1:300 dilution, Goat anti-Mouse IgG (H+L) Alexa Fluor 488 (A11001, Invitrogen) at 1:200 dilution, anti-NQO1 (AB34173, Abcam) at 1:200 dilution, anti-NQO1 (11451-1-AP, Proteintech) at 1:200 dilution, anti-G3BP1 (AB56574, Abcam) at 1:200 dilution, anti-G3BP1 (sc-365338, Santa Cruz) at 1:200 dilution, anti-G3BP1 (13057-2-AP, Proteintech) at 1:200 dilution, anti-DCP1A (SC-100706, Santa Cruz) at 1:200 dilution, and anti-DYKDDDDK-tag (20543-1-AP, Proteintech) at 1:2000 dilution.

The human pancreatic cancer tissue microarray (Ethics Approval Number-SHYJS-CP-2210045) was obtained from Shanghai Xinchao Biotechnology Co., Ltd. (Shanghai, China). Immunofluorescence staining was performed following standard protocols previously established for murine tissues. Primary antibodies included rabbit anti-NQO1 (AB34173, Abcam; 1:200 dilution) and mouse anti-G3BP (AB56574, Abcam; 1:500 dilution). Secondary antibodies were incubated for 1 h at room temperature: Goat anti-Rabbit IgG (H+L) conjugated with Alexa Fluor 488 (A-11008, Invitrogen; 1:200) and Goat anti-Mouse IgG (H+L) labeled with Alexa Fluor 594 (A-11005, Invitrogen; 1:200).

### Co-Immunoprecipitation

The 6-well plates were pre-transfected with the target protein plasmids (n=2). After a 48-hour incubation period, the cells were harvested, and the contents from two wells were pooled into a single microcentrifuge tube. Following washing with PBS, Complete Lysis Buffer for Western and IP (P0037, Beyotime Biotechnology) supplemented with a protease inhibitor cocktail (HY-K0010, MCE) was added. The samples were rotated at 4°C for 30 minutes and then centrifuged at 20,000 × g for 20 minutes at 4°C. Anti-HA magnetic beads (88836, Gibco) or anti-DYKDDDDK magnetic beads (A36797, Thermo Fisher) were pre-treated with TBST and NETE buffer, using 2 μl per test. The supernatant from the centrifuged protein lysate was collected, and 40 μl aliquots were mixed with 10 μl of 5×loading buffer as the input sample, followed by heating at 95°C for 5 minutes. The remaining supernatant was incubated with the pre-treated magnetic beads at 4°C for 2 hours under continuous rotation.

The magnetic beads were sequentially washed: first with TBST buffer, then with NETE buffer, and finally with NETN buffer containing 2% Tween-20. Each washing step was repeated twice to ensure the complete removal of non-specific binding components. Protein elution was performed by incubating the beads with 45 μl of Complete Lysis Buffer for Western and IP supplemented with a protease inhibitor cocktail and 1× loading buffer at 95°C for 5 minutes.

### Western blot

Cellular lysis was performed using RIPA buffer (150 mM sodium chloride, 50 mM Tris-HCl pH 7.4, 0.1% SDS, 1 mM EDTA, 1% NP-40, 1% sodium deoxycholate) supplemented with a protease inhibitor cocktail. Total protein extracts were then subjected to SDS-PAGE and subsequently transferred onto PVDF membranes. After blocking with 5% non-fat milk for 1 hour at room temperature, the membranes were incubated with primary antibodies at 4°C overnight, including anti-NQO1 (1451-1-AP, Proteintech) at a dilution of 1:3000, anti-G3BP1 (13057-2-AP, Proteintech) at 1:10000, anti-β-Actin (66009-1-Ig, Proteintech) at 1:50000, anti-DYKDDDDK-tag (20543-1-AP, Proteintech) at 1:20000, anti-DYKDDDDK-tag (66008-4-Ig, Proteintech) at 1:20000, anti-HA (51064-1-AP, Proteintech) at 1:8000, anti-TIA1 (12133-2-AP, Proteintech) at 1:1000, and anti-HSP90 (13171-1-AP, Proteintech) at 1:5000. The membranes were then washed three times with TBST (10 minutes per wash) at room temperature, followed by incubation with HRP-conjugated secondary antibodies (SA00001-1 and SA00001-2, Proteintech) for 1 hour. After an additional three washes with TBST (10 minutes per wash) at room temperature, protein bands were visualized using the Tanon-5200 chemiluminescence imaging system with ECL substrate, and the resulting images were analyzed and quantified using ImageJ software.

### Protein purification, staining and in vitro fluorescence-based droplet formation assay

The pGEX-4T-1-NQO1-GST plasmid and pET-28a-G3BP1-6 × His plasmid were transformed into T7 Express competent cells and Rosetta DE3 competent cells (purchased from Weidibio), respectively. Protein expression was induced by the addition of 0.6 mM IPTG (T1416-5G, Beyotime) and maintained at 16°C for overnight incubation, subsequent to which bacterial lysis was performed utilizing Hammer super germ solution (BR0005-02, ACE). The lysate supernatant was incubated with glutathione-agarose beads (P2262, Beyotime) or Ni-NTA resin (P2226, Beyotime) at 4°C overnight. Following the washing procedure, glutathione-agarose beads and Ni-NTA resin were subjected to either imidazole gradient elution or competitive elution with glutathione (GSH), respectively. Through multistep chromatographic purification, highly purified GST-NQO1 and His-G3BP1 proteins were obtained, which were subsequently eluted into sodium bicarbonate (NaHCO_3_, pH=8.3) buffer for next protein lable experiment.

The GST-NQO1 and His-G3BP1 proteins were specifically labeled with AF488 NHS ester (40779ES03, Yeasen Biotechnology) through a 1 hour incubation process at room temperature. (The molar ratio of AF488 NHS ester to target protein was precisely maintained at 2:1, with the reaction conducted at pH 8.3). Following the incubation period, the labeling reaction was effectively terminated by the addition of 200 mM Tris buffer (pH 8.0), while simultaneously quenching any unbound dye molecules. The fluorescently labeled proteins were subsequently purified into a buffer solution containing 50 mM Tris-HCl and 150 mM NaCl (pH 7.5) using a HiTrap desalting column. The purified protein was then analyzed using a multichannel live cell imaging system after being combined with either Ficoll400 molecular biology reagent (F2637, Sigma) (to replicate intracellular crowding conditions), or total RNA, respectively.

### Recovery Experiment Following Intracellular Fluorescence Photobleaching

Mia-PaCa2 cells stably overexpressing EGFP-NQO1 protein through lentiviral transduction were seeded into confocal imaging dishes. Photobleaching was induced using a 488 nm laser beam in the confocal system (Leica STELLARIS 5, Germany), with a defined bleaching area of 2 μm. Fluorescence recovery was monitored and recorded at 2-second intervals.

### GST pull-down assay

The purified GST-NQO1 and His-G3BP1 proteins were reconstituted in TBS buffer (25 mM Tris-HCl, 150 mM NaCl, pH 7.2) and subsequently concentrated to a final concentration of ≥ 1 μg/μL. GSH agarose beads (21516, Thermo Fisher Scientific) underwent five washing cycles using a wash solution (pull-down lysis buffer: TBS buffer=1:1). Subsequently, 150 μg of the bait protein, GST-NQO1/GST, was immobilized onto the beads at 4°C for 2 hours, followed by five additional washing steps. The prey protein, His-G3BP1 (150 μg), was then incubated with the immobilized bait protein at 4°C overnight, after which the beads were washed five times. For each experimental sample, 250 μl of elution buffer (10 mM GSH in TBS buffer) was applied, and the eluate was collected by centrifugation at 1,250 g for 1 minute following gentle agitation for 5 minutes. The GST-NQO1/GST and His-G3BP1 proteins were mixed at a 1:1 mass ratio as the input sample. Both the input and elution fractions were subsequently analyzed by SDS-PAGE and coomassie brilliant blue staining.

### ROS Quantification and Analysis

Mia-PaCa2 Vector and NQO1 KO cells were cultured in 6-well plates containing pre-mounted slides 24 hours prior to the experiment. Following the protocol of the ROS detection kit (S0033S, Beyotime), the culture medium was replaced with serum-free medium containing DCFH-DA (10 μM) at a 1:1000 dilution and ROS positive control at a 1:100 dilution. The cells were subsequently incubated at 37°C for 20 minutes in a humidified cell culture incubator. To ensure complete removal of extracellular DCFH-DA, the cells were washed three times with serum-free medium. In accordance with the experimental grouping design, NAC (5 mM) (S0077, Beyotime) was initially administered and incubated for 1 hour at 37°C, followed by the addition of NaAsO_2_ (1 mM) for an additional 1 hour at 37°C. The slides were then processed for immunofluorescence analysis. Concurrently, the remaining cells in the 6-well plates were trypsinized, resuspended in serum-free medium, and filtered through a 200-mesh double-layer filter. The cell suspensions were subsequently transferred to flow cytometry tubes for further analysis. The FITC (Fluorescein Isothiocyanate) fluorescence intensity, indicative of intracellular ROS levels, was quantitatively assessed utilizing Flow Cytometry Fortessa (BD LSRFortessa, USA).

### Cellular proliferation analysis

24 hours prior to the experimental procedure, cells were cultured in 96-well culture plates. Following a 48 hours transfection period with either plasmid or siRNA, 10 μl of CCK-8 working solution (C0038, Beyotime) was administered to each well at consistent time intervals daily, followed by incubation at 37°C for 60 minutes. Cell viability was quantitatively assessed at an absorbance wavelength of 450 nm utilizing a Cytation 3 Cell Imaging Multimodal Reader (BioTek, USA).

### Mice

*Pdx1^Cre^;* LSL*-Kras*^G12D^; *Nqo1^-/-^*(KCN^-/-^) mice were generated through crossbreeding of the following strains: *Pdx1-Cre* (J-014647), LSL*-Kras^G12D^* (Cat.NO. NM-KL-190003) and *NQO1^-/-^*(Cat. NO. NM-KO-190845) (Shanghai Model Organisms Center, Inc, 2022). As control groups*, Pdx1^Cre^;* LSL*-Kras*^G12D^ (KC) mice were produced by crossing *Pdx1^Cre^* and LSL*-Kras*^G12D^ strains. All mice were of the C57BL/6J strain and maintained at a specific pathogen-free (SPF) grade. The rearing conditions were controlled as follows: room temperature of 23 ± 1°C, relative humidity ranging from 40% to 70%, animal illumination at 20 lx, a light-dark cycle of 12 h/12 h, and ad libitum access to food and water. All experimental procedures were performed in strict compliance with national ethical guidelines for animal research and were formally approved by the Animal Ethics Committee of Xuzhou Medical University (Approval No. 202410T003).

Pancreatic tissues from 3-month-old and 6-month-old KC and KCN^-/-^ mice were collected, with approximately 30 mg of tissue homogenized in RIPA buffer containing protease and phosphatase inhibitors, and approximately 20 mg preserved in RNA later solution and subsequently fragmented. The remaining pancreatic tissue was processed as follows: overnight fixation in 4% paraformaldehyde (PFA), followed by sequential dehydration through graded ethanol series (50%, 75%, 85%, 95% I, 95% II, 100% I, and 100% II) with 1-hour intervals for each concentration. Subsequently, the tissues were cleared in xylene I and II (10 minutes each), infiltrated with paraffin I and II (1 hour each), and finally immersed in paraffin III overnight. The following day, the tissues were embedded in paraffin blocks. Serial sections of 0.35 μm thickness were prepared and submitted to Wuhan Servicebio Technology for comprehensive histopathological analysis, including hematoxylin and eosin (H&E) staining, Amylase and CK19 immunofluorescence staining, Sirius red staining, Alcian blue staining, Ki67 immunohistochemistry, and G3BP immunofluorescence staining.

For histological analysis, tissue sections were subjected to Alcian blue, Sirius red, and H&E staining procedures, involving sequential steps of dye application, dehydration, and mounting. Regarding Ki67 immunohistochemical staining, paraffin-embedded sections underwent initial dewaxing and rehydration, followed by antigen retrieval. Endogenous peroxidase activity was subsequently blocked using 3% hydrogen peroxide. After a 30-minute blocking step with 3% bovine serum albumin (BSA), the primary antibody was applied and incubated overnight at 4°C. The secondary antibody incubation was performed at room temperature for 50 minutes, followed by 3,3’-diaminobenzidine (DAB) chromogenic development. Finally, nuclear counterstaining was achieved using hematoxylin, and the sections were dehydrated and mounted for microscopic examination.

Regarding immunofluorescence staining of paraffin sections, the preparation process involved deparaffinization and rehydration of sections, antigen retrieval, and blocking with BSA. Primary antibody incubation was conducted at 4°C overnight, followed by secondary antibody incubation at room temperature for 50 minutes. Additionally, DAPI nuclear counterstaining and tissue autofluorescence quenching were performed prior to slide mounting.

The following antibodies were utilized: G3BP (AB56574, Abcam) at a dilution of 1:500, Cy3-conjugated goat anti-mouse IgG (GB21301, Servicebio) at 1:300, Anti-CK19 (AB52625, Abcam) at 1:200, Anti-Amylase (SC-46657, Santa Cruz) at 1:200, Cy3-labeled goat anti-rabbit IgG (GB21303, Servicebio) at 1:300, Alexa Fluor 488-labeled goat anti-mouse IgG (GB25301, Servicebio) at 1:400, Anti-Ki67 (Servicebio, GB111141) at 1:1000, and HRP-labeled goat anti-rabbit secondary antibody (Servicebio, GB23303) at 1:200.

### Three-Dimensional Organoid Culture System Establishment

Prior to the experimental procedure, a 24-well plate was prepared by coating each well with 250μl of 85% type I collagen solution (composed of 85% rat type I collagen, 1M NaOH, and 10% 10 × PBS in ddH_2_O) as the basal layer, which was subsequently polymerized in a 37°C incubator for 4 hours. Pancreatic tissues were excised from four 8-to 10-week-old KC or KCN^-/-^ mice, with two mice representing each genotype. Each pancreatic specimen underwent enzymatic digestion using 0.5-0.75 ml of 2 mg/ml collagenase P (in 1×HBSS) at 225 rpm and 37°C for 20-30 minutes, with the supernatant condition being assessed at 10-minute intervals. Upon detection of supernatant turbidity, the digestion process was immediately terminated by the addition of 5 ml of pre-chilled 1× HBSS containing 5% FBS. The samples were then subjected to centrifugation at 2000 rpm for 2 minutes at 4°C, followed by supernatant removal and multiple washing cycles (minimum of two repetitions). After the final wash, the cellular pellet was reconstituted in 5 ml of 1× HBSS supplemented with 5% FBS. The resultant suspension was filtered through a 100 μm strainer and gently layered along the wall into 20 ml of pre-cooled HBSS containing 30% FBS for density gradient separation. Following centrifugation at 1000 rpm for 2 minutes, the supernatant was carefully discarded, and the pellet was resuspended in 1-2 ml of organoid medium (RPMI 1640 supplemented with 1% FBS, 1% penicillin-streptomycin, 0.1 mg/ml STI, and 1μg/ml dexamethasone). Subsequently, 100 μl of cell suspension (containing 50,000-100,000 cells) was thoroughly homogenized with 250 μl of 85% type I collagen and meticulously layered along the wall of the 24-well plate as the intermediate layer, followed by incubation at 37°C for 4 hours. Finally, 300-400 μl of organoid medium was added to each well as the top layer. The following day, acinar cells were inoculated with the viral solution. The experimental analysis was performed on the fourth day.

### Quantification and statistical analysis

Data analysis and statistical analysis was performed using GraphPad Prism v9.3.1. All data are represented as mean±s.d. The significance of intergroup differences was assessed using two-tailed unpaired Student’s t-tests. The statistical parameters are reported in the graphs and the corresponding legends. Statistical differences among multiple groups in Fig. 5a were analyzed by Kruskal-Wallis test, and pairwise differences between groups were further examined using the Wilcoxon rank sum test. The obtained *p*-values are annotated in the figure, with *p* < 0.05 considered statistically significant. In Fig. 6f, correlation analysis was performed using Pearson test. Pearson’s correlation coefficients are shown in the figure, with R values less than 0.3 considered as weak correlations. The sample sizes (number of cells analyzed) for each experimental condition are explicitly indicated in the respective figure legends.

## Notes

### Competing Interest Statement

The authors have declared no competing interest.

