## Supplementary figures and images for "NQO1 phase condensation promotes stress granule assembly to facilitate pancreatic carcinogenesis"

### Supplemental Figure 1

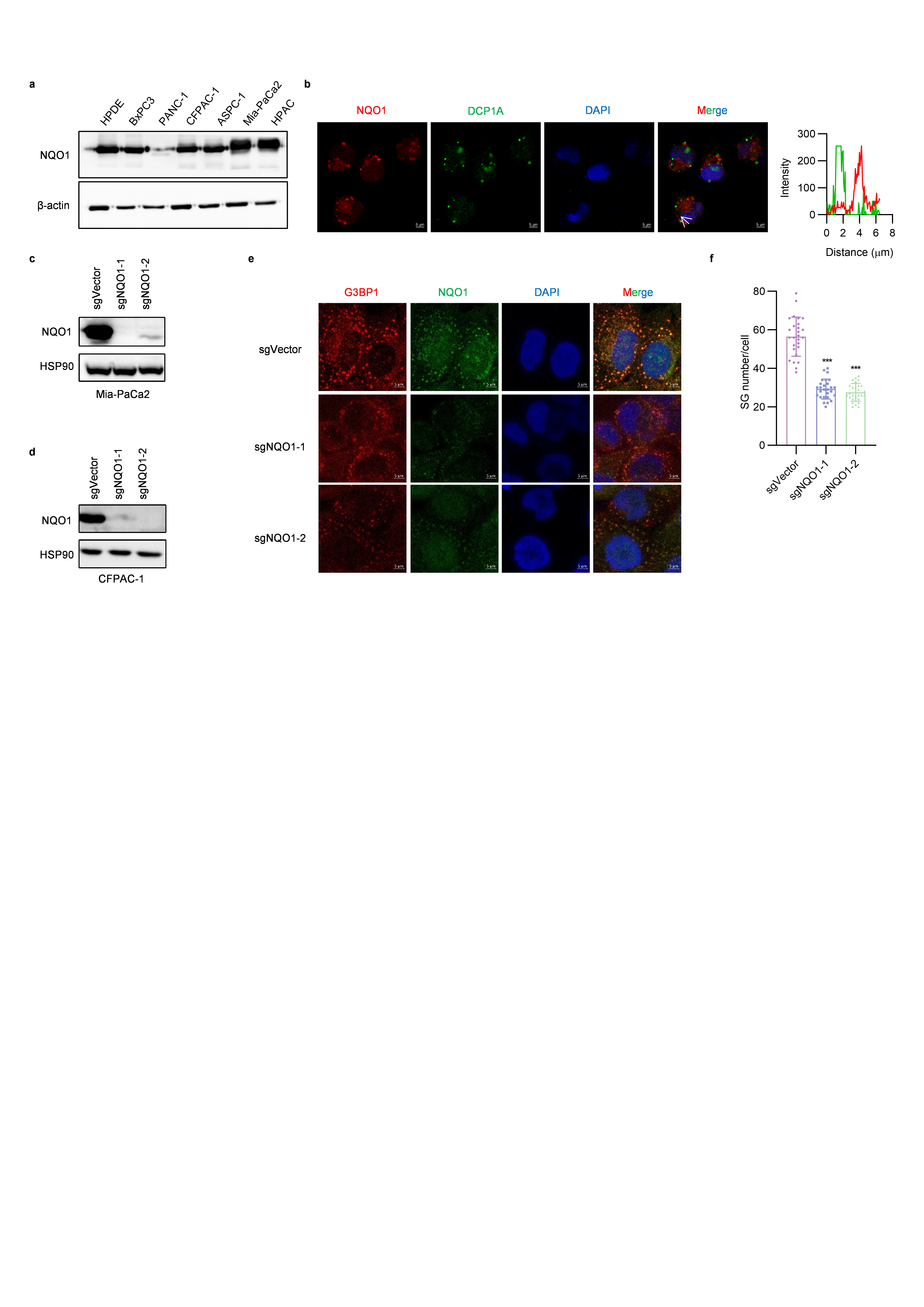

### Supplemental Figure 2

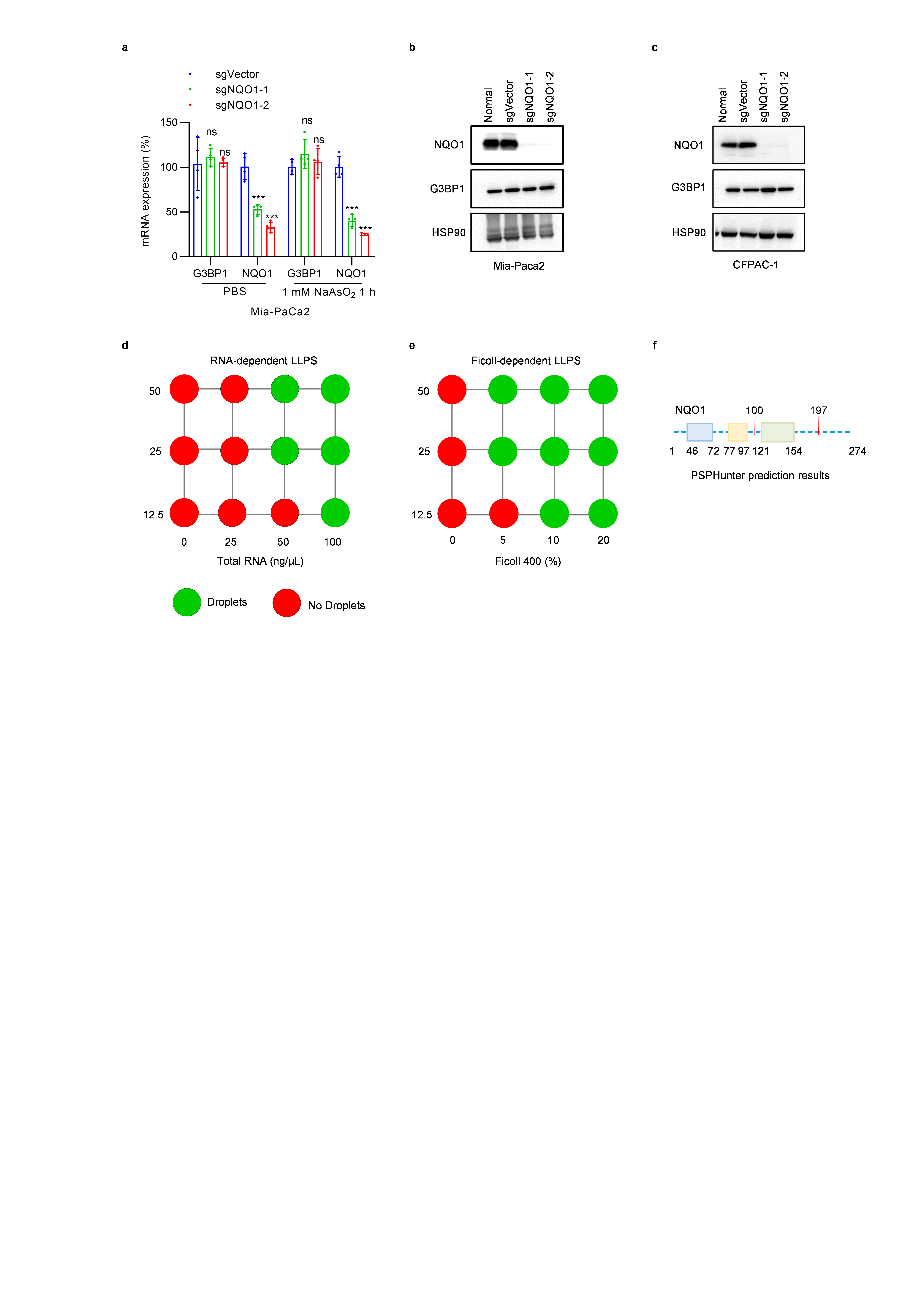

### Supplemental Figure 3

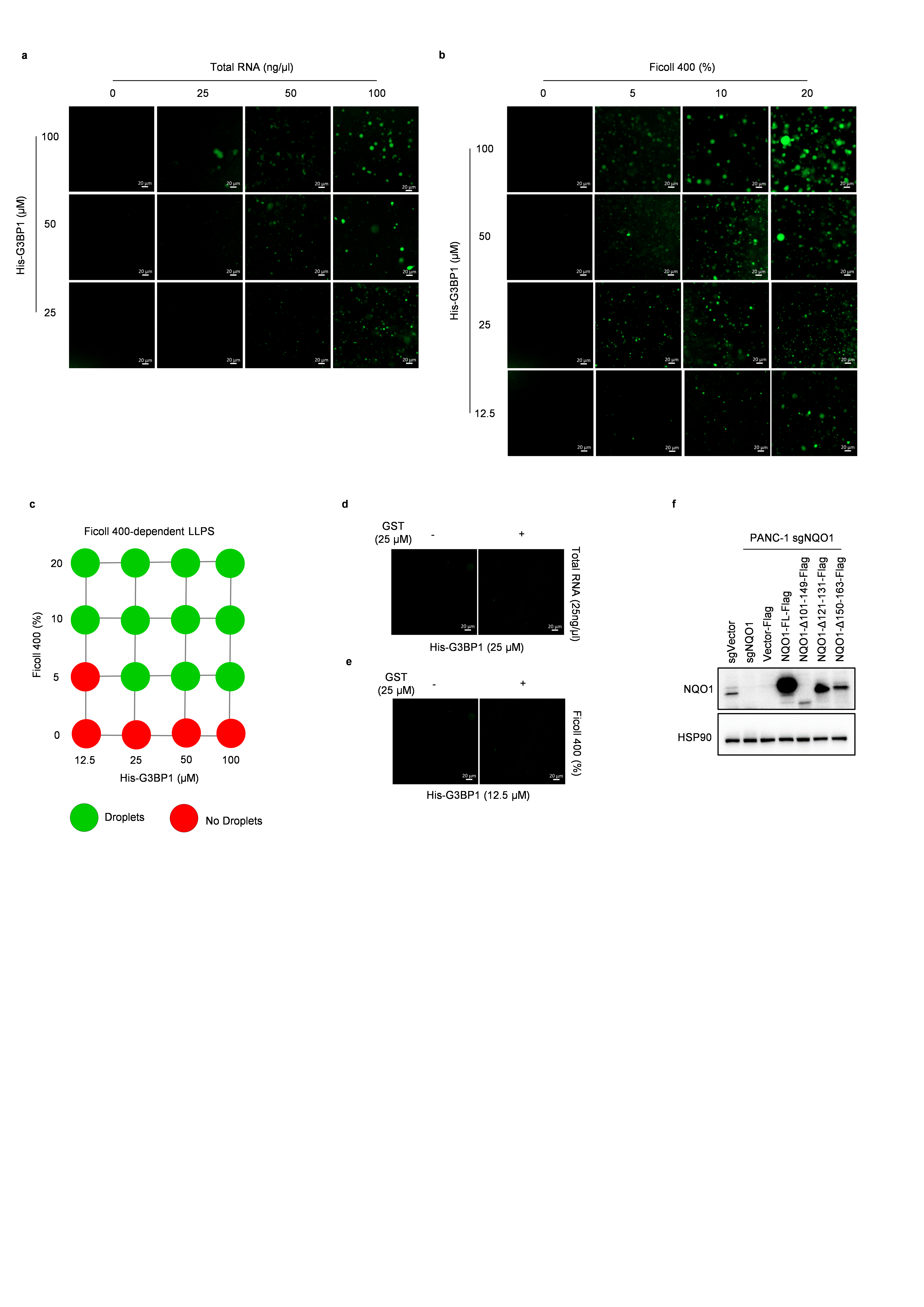

### Supplemental Figure 4

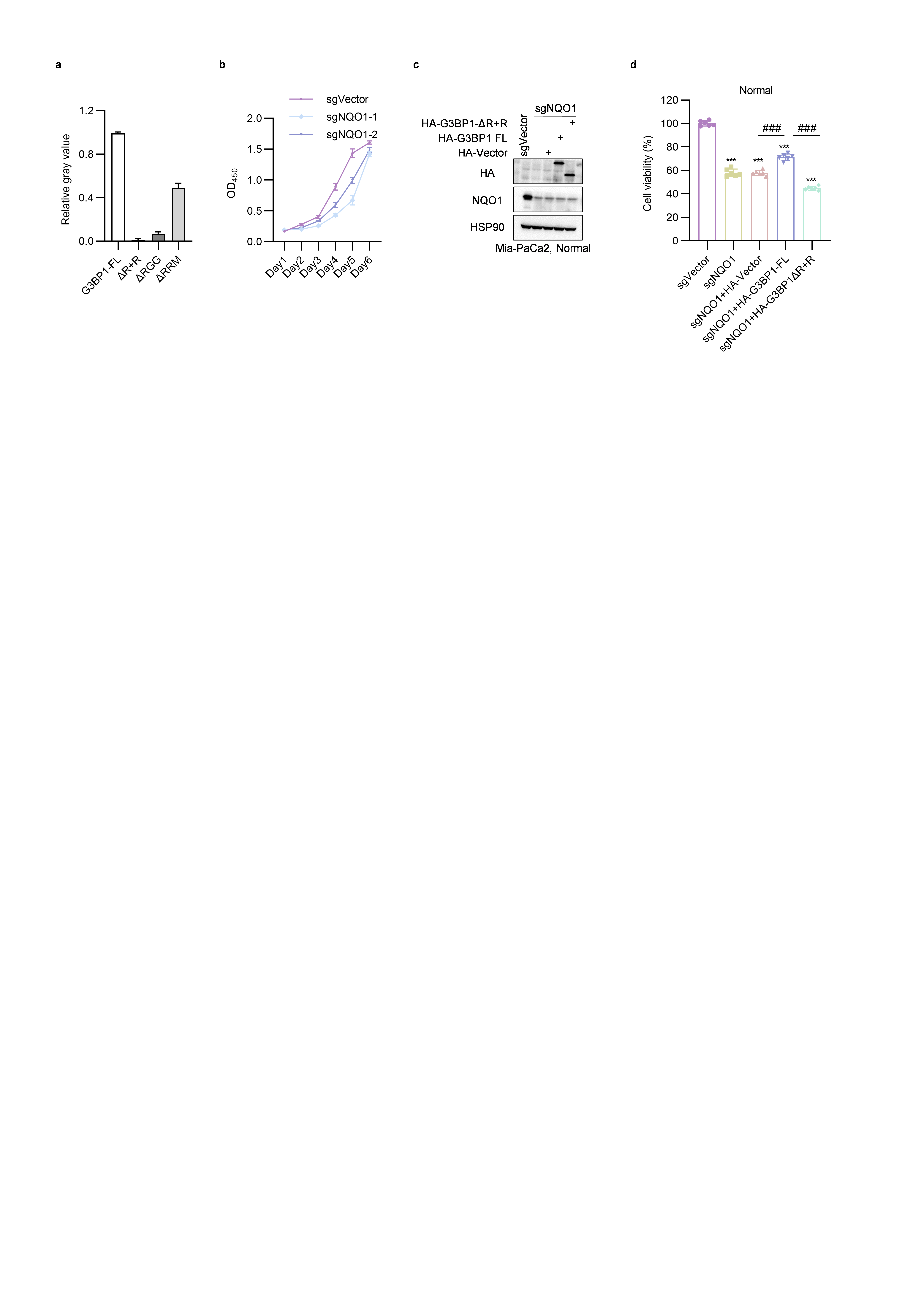

### Supplemental Figure 5

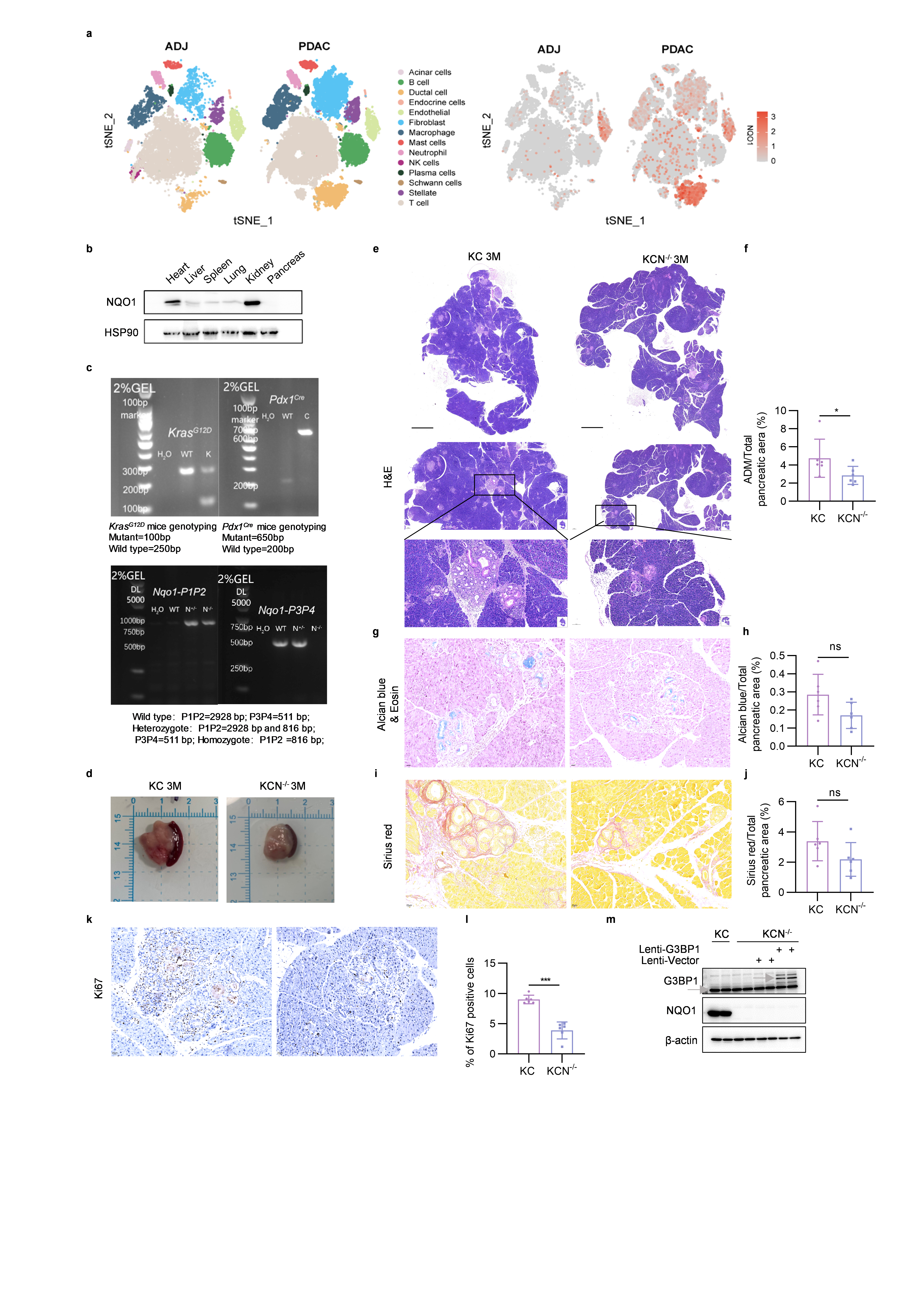
